# *Cel*EsT: a unified gene regulatory network for estimating transcription factor activities in *C. elegans*

**DOI:** 10.1101/2024.06.26.597625

**Authors:** Marcos Francisco Perez

## Abstract

Transcription factors (TFs) play a pivotal role in orchestrating the intricate patterns of gene regulation critical for development and health. Although gene expression is complex, differential expression of many genes is often due to regulation by just a handful of TFs. Despite extensive efforts to elucidate TF-target regulatory relationships in *C. elegans*, existing experimental datasets cover distinct subsets of TFs and leave data integration challenging.

Here I introduce *Cel*EsT, a unified gene regulatory network (GRN) designed to estimate the activity of 487 distinct *C. elegans* TFs - ∼58% of the total - from gene expression data. To integrate data from ChIP-seq, DNA-binding motifs, and eY1H screens, different GRNs were benchmarked against a comprehensive set of TF perturbation RNA-seq experiments and identified optimal processing of each data type. Moreover, I showcase how leveraging conservation of TF binding motifs in the promoters of candidate target orthologues across genomes of closely-related species can distil targets into a select set of highly informative interactions, a strategy which can be applied to many model organisms. Combined analyses of multiple datasets from commonly-studied conditions including heat shock, bacterial infection and male-vs-female comparison validates *Cel*EsT’s performance and highlights previously overlooked TFs that likely play major roles in co-ordinating the transcriptional response to these conditions.

*Cel*EsT can be used to infer TF activity on a standard laptop computer within minutes. Furthermore, an *R Shiny* app is provided for the community to perform rapid analysis with minimal coding experience required. I anticipate that widespread adoption of *Cel*EsT will significantly enhance the interpretive power of transcriptomic experiments, both present and retrospective, thereby advancing our understanding of gene regulation in *C. elegans* and beyond.

## INTRODUCTION

Transcription factors (TFs) have a central role in co-ordinating development, the maintenance of cell identity and in transcriptional responses to insults. TFs act by binding to specific sites in the genome, recruiting regulatory proteins, transcriptional machinery or chromatin modifiers to influence the expression of their nearby target genes (Lambert *et al*. 2018). While in humans the targets of TFs may be distantly located along the chromosome due to phenomena including chromatin looping (Palstra and Grosveld 2012), the popular model organism *Caenorhabditis elegans* exhibits a simpler regulatory model whereby most regulation occurs by TFs binding to the promoter region of a gene proximal to the transcription start site (Reinke *et al*. 2013).

Due to their essential roles, TFs are strongly conserved across long evolutionary timescales. This has allowed model organisms like *C. elegans* to play a key role in the discovery and characterisation of the effector TFs downstream of many pathways that are common in mammalian development. For example, the SMAD TFs that co-ordinate transcription downstream of the TGF-beta signalling pathway were first described and named for their mutant phenotypes in *C. elegans* and *Drosophila melanogaster* (Padgett *et al*. 1998). Interestingly, the preferred DNA sequence motifs of TFs are also often strongly conserved even when overall protein sequence similarity is low (Lambert *et al*. 2019). For example, after >500 million years of divergence, the human and *C. elegans* orthologues of the transcription factor TFEB/HLH-30 are just ∼44% similar but their core DNA binding motif has remained essentially constant.

A host of post-transcriptional mechanisms exist to modify TFs in ways that strongly influence their gene regulatory activity (Reinke *et al*. 2013). As such, the expression of transcription factors (at the mRNA or protein level) often does not reflect their activity. For example, the FOXO orthologue DAF-16 is phosphorylated downstream of insulin signalling, which leads to its exclusion from the nucleus and consequent inhibition of regulatory activity (Cahill *et al*. 2001; Lin *et al*. 2001). Post-translational control of TF activity is particularly critical when environmental insults necessitate a rapid transcriptional response, such as in the case of the activation of HSF-1 by heat shock (Vihervaara and Sistonen 2014). However, although changes in TF activity can sometimes be inferred by visualising changes in subcellular localisation or profiling genomic binding, they are only directly detectable at the transcriptional level by the concerted effect of altered TF activity on their various target genes.

Several existing methods can provide quantitative estimates of TF activity from gene expression data based on the expression of their target genes (Badia-i-Mompel *et al*. 2022). However such approaches also require prior knowledge in the form of a gene regulatory network (GRN) that lists which genes are regulated by each particular TF (also referred to as a TF’s regulon; (Garcia-Alonso *et al*. 2019)). While great effort has been expended defining TF regulons in *C. elegans* via different experimental methods (Narasimhan *et al*. 2015; Fuxman Bass *et al*. 2016; Kudron *et al*. 2018; Kudron *et al*. 2024), each individual dataset covers a distinct subset of TFs and integration of the disparate data available has been challenging. As a result, the *C. elegans* community has to date lacked a unified TF-target prior knowledge resource to enable TF activity estimation and unlock the immense mechanistic insights it can provide.

The aim of this study is to integrate distinct available genome-wide resources on TF-target interactions into a unified *C. elegans* GRN which could be used with existing methods for quantitative TF activity estimation. A comprehensive benchmarking dataset of TF perturbation RNA-seq experiments was compiled with which to quantitatively assess the performance of different data sources, data processing strategies and methods for TF activity estimation. Applied to a validation dataset of RNA-seq experiments from various genetic, environmental or physiological conditions, the resulting GRN, christened *Cel*EsT, recapitulated known TF biology and also highlighted the potential importance of previously overlooked TFs. TF activity estimation with *Cel*EsT will provide *C. elegans* researchers with powerful mechanistic insights into transcriptomic experiments via rapid analysis on local computers using *R*, *Python* or a dedicated *R Shiny* app made available to the community at https://github.com/IBMB-MFP/CelEsT-app.

## RESULTS

Using transcriptomic data and a prior-knowledge GRN detailing TF-target relationships, TF activity can be estimated via several existing methods (available through the *decoupler* software package (Badia-i-Mompel *et al*. 2022)). Of the methods which gauge activity by fitting a linear model to target gene expression, the univariate linear model (‘ulm’) method considers TFs separately, whereas the multivariate linear model (‘mlm’) method fits a single model to all TFs. The latter can thereby theoretically disentangle the separate effects of distinct TFs with overlapping targets. The weighted sum method (‘wsum’) involves summing scores for all targets of a given TF. The *decoupler* package can also provide a consensus score which integrates the scores from disparate methods.

### Three large-scale experimental datasets report TF-target interactions in *C. elegans*

To construct GRNs for *C. elegans*, 3 major resources were identified for selection of TF regulatory targets, originating from distinct experimental techniques (Fig 1A). The first was modERN (Kudron *et al*. 2018; Kudron *et al*. 2024); heir to the modENCODE project (Gerstein *et al*. 2010), consisting of 525 ChIP-seq experiments for 354 TFs. The second was CisBP (Narasimhan *et al*. 2015), a resource compiling DNA binding motifs for 252 motifs identified largely by *in vitro* methods, either determined directly or inferred by high similarity of a TF’s DNA-binding domain to other TFs with characterised binding sequence specificity (Lambert *et al*. 2019). The third data resources was a collection of TF-gene promoter interactions identified from enhanced yeast one-hybrid assays (eY1H) (Fuxman Bass *et al*. 2016). In addition to ChIP-seq experiments from modERN, ChIP-seq datasets were identified for another 5 TFs not included in modERN from the NCBI Gene Expression Omnibus (GEO). In total information was recovered to identify potential targets for 596/833 *C. elegans* TFs (Kudron *et al*. 2024). Additional sources for TF-target interactions were considered but ultimately discarded (see Methods). The networks resulting from this study are unsigned i.e. they do not assign a mode of regulation (e.g. activator or repressor) to each TF. Nonetheless TF modes of regulation compiled from WormBase and UniProt can be found in Table S1 and in the output of the *Cel*EsT *Shiny* app.

**Fig 1.**
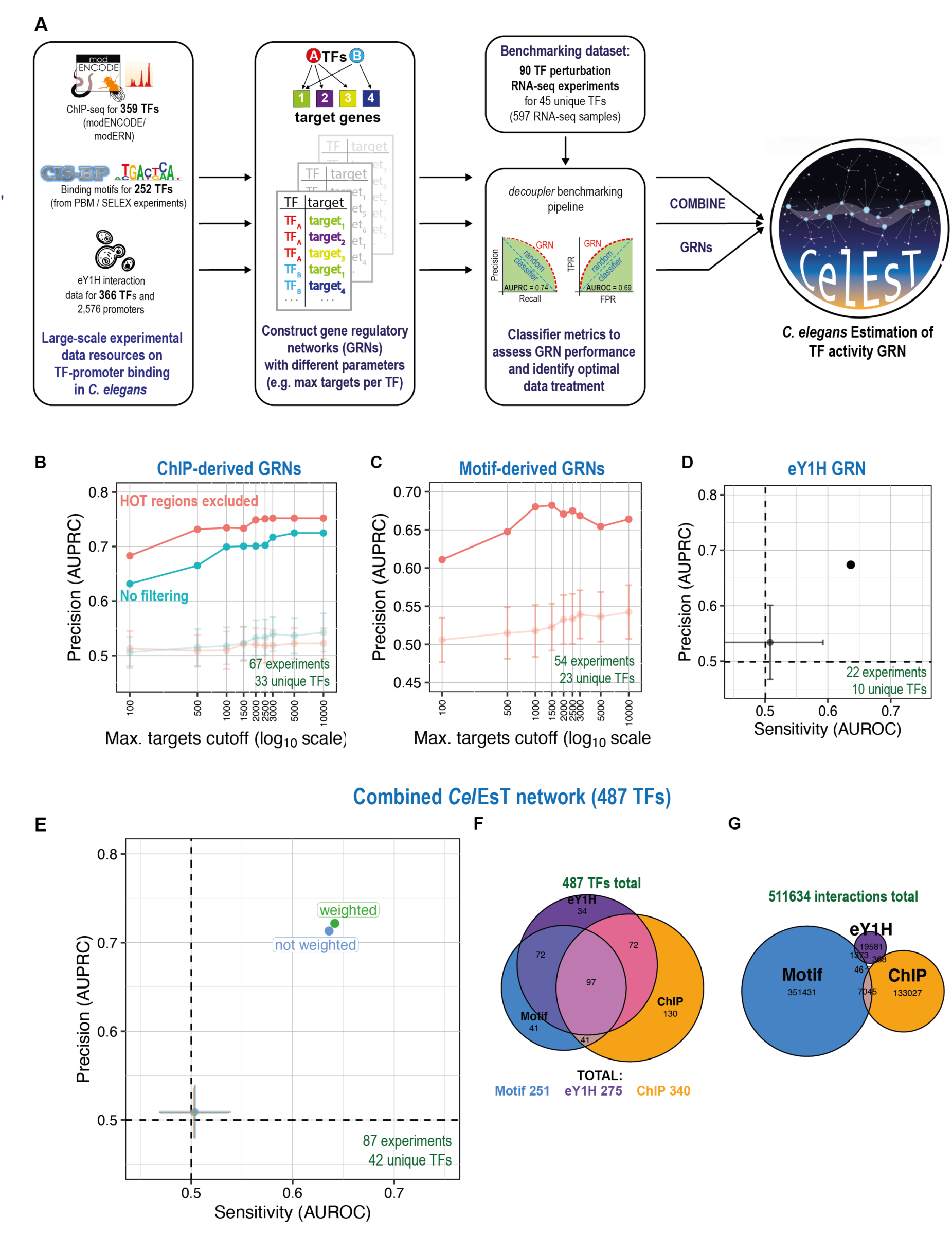
Generation and benchmarking of GRNs from publicly available large-scale datasets. **A)** *Cel*EsT synthesises TF-target information from 3 large-scale and distinct experimental sources: TF ChIP-seq experiments from the modERN consortium, TF DNA-binding motifs directly measured or inferred from PBM/SELEX experiments from the CisBP database and a dataset of enhanced yeast one-hybrid (eY1H) assays measuring direct TF-promoter interactions. GRNs are constructed and assessed against a benchmarking pipeline using the *decoupler* software package. Optimised networks are combined to yield a unified network. **B)** Line shows evolution of AUPRC in benchmarking pipeline with increasing cut-off for maximum number of targets per TF for GRNs derived from ChIP-seq data with (red) or without (blue) filtering of HOT regions. Note x-axis (cut-off) is on a log scale. Multivariate linear model (‘mlm’) method is shown. Faded lines show AUPRC mean and standard deviation (SD) for randomised networks (100 iterations). See also Fig S1B, C, E & F for performance with other methods. **C)** Line shows evolution of AUPRC in benchmarking pipeline for the ‘mlm’ method with increasing cut-off for maximum number of targets per TF (log scale) for GRNs derived from TF DNA binding motifs. Faded line shows AUPRC mean/SD for 100 randomised networks. See also Fig S2B-C for performance with other methods. **D)** Benchmarking performance (AUPRC and AUROC) for a GRN with regulons derived from TF-promoter interactions demonstrated in an eY1H assay. Lighter point with error bars shows mean/SD for randomised network. See also Fig S3A for other methods. **E)** Benchmarking performance for the *Cel*EsT network, which covers 487 TFs by combining data from multiple experimental sources. See also Fig S3C for other methods. **F)** Euler diagram showing contribution of TFs to the *Cel*EsT network from the three distinct experimental datasets. Total unique TFs shown above, total from each dataset shown below. **G)** Euler diagram showing contribution of TF-target interactions to the *Cel*EsT network from the three distinct experimental datasets. Total TF-target interactions shown above. See also Fig S3D for interactions considering only TFs common to all three datasets.

In order to judge the performance of the GRNs resulting from the integration of these data resources, I used the benchmarking pipeline from the *decoupler* package (Fig 1A) (Badia-i-Mompel *et al*. 2022). This pipeline assesses the ability of TF activity estimations to correctly identify perturbed TFs from expression of their target genes in a benchmarking dataset. This allows for a quantitative performance comparison both for different GRNs and for different statistical methods of estimating TF activity, using the classifier metrics AUROC and AUPRC. Although AUROC/AUPRC are related and correlated metrics, they can display opposite trends (e.g. Fig S1C) and so GRNs that maximised both AUROC and AUPRC were prioritised where possible. As a benchmarking resource, 90 *C. elegans* RNA-seq experiments were identified from the Gene Expression Omnibus (GEO) repository with TF loss of function (e.g. by mutation or RNAi knockdown) or gain-of-function (via overexpression or gain-of-function alleles) with matching controls for 45 unique TFs from 73 distinct studies (Table S2). RNA-seq datasets were uniformly processed and differential expression (DE) analysis was performed between TF perturbation and control samples, including accounting for any potential mismatch in developmental stage resulting from the knockdown ((Bulteau and Francesconi 2022); see Methods). The collected DE stats for all studies were used as the benchmarking dataset.

### Excluding targets in HOT regions improves performance of ChIP-derived GRNs

Potential TF targets from ChIP-seq experiments were identified by TF binding to the promoter. Targets were prioritised firstly by recurrence over multiple experiments, if different life stages had been assayed for the same TF (∼20% of modERN TFs), and secondly by strength of ChIP peak signal. Given that some ChIP peaks might represent false positives, the benchmarking pipeline was used to identify a suitable cut-off for maximum number of targets per TF (Fig S1A). However I found that the best performance resulted from applying no cut-off to ChIP-derived regulons (Fig 1B, Fig S1B, C, E & F).

It has been reported from large-scale ChIP-seq datasets that some genomic sites appear to be promiscuously bound by large numbers of TFs - so-called High Occupancy Target (HOT) regions (Kvon *et al*. 2012). Including targets in HOT regions with many bound TFs could potentially complicate the attribution of expression differences at these loci to activity of specific TFs. To address this, gene promoters within HOT regions were excluded (Fig S1D), substantially improving benchmarking pipeline performance at all cut-offs and for all TF activity estimation methods (Fig S1B, C, E & F), though leading to the loss of 44 TFs that retained little or no promoter binding outside of HOT regions.

Of note, comparison between TF activity estimation methods using ChIP-derived GRNs demonstrated that the multivariate linear model (‘mlm’) method had both the best overall performance and the best discrimination versus randomised networks, with substantially worse and near-identical performance for the univariate linear model (‘ulm’) and weighted sum (‘wsum’) methods (Fig S1B-F).

### Optimal regulon size cut-offs are critical for GRNs derived from TF DNA-binding motifs

Gene promoter sequences were scanned for significant matches to *in vitro*-derived DNA binding motifs to identify potential targets for 252 TFs (Fig S2A). In contrast to ChIP-derived GRNs, benchmark performance clearly peaked at a maximum cut-off of 1500 targets for each TF (Fig 1C, Fig S2B-C). While the ‘mlm’ method was still the best performing individual method, unlike the ChIP-based GRNs the motif-based GRN also performed well with both ‘ulm’ and ‘wsum’ (Fig S2B-C).

Regulatory regions may have several distinct binding sites for the same TF (Kazemian *et al*. 2013) which can lead to high TF occupancy despite lower affinity of individual sites (Crocker *et al*. 2016; Shahein *et al*. 2022), a phenomenon which has been called homotypic binding site clustering (Payne and Wagner 2015) (Fig S2D). I hypothesised that taking into account the presence of multiple binding sites for a given TF, rather than simply taking the best match, might improve network performance. To reflect the possibility of homotypic binding, a combined score was computed providing additional but diminishing points for multiple significant binding sites in the candidate target gene’s promoter region. The effect of this is to change the ordering of potential target genes and so potentially the composition of genes are included within a given cut-off. While this combined homotypic binding score did improve performance using very low cut-offs, at the intermediate cut-offs previously identified as optimal it had the opposite effect (Fig S2E). As such, I proceeded using GRNs compiled from target lists ordered using only the strength of the best sequence match to the interrogated motif.

Additionally a GRN was compiled from 160 TFs with at least 15 target promoters from the eY1H TF-promoter interaction dataset in (Fuxman Bass *et al*. 2016). This GRN displayed good performance in the benchmarking dataset, thus clearly contributing valuable regulatory information (Fig 1D, Fig S3A).

### Integrated GRNs from multiple data sources have improved performance

The optimal ChIP- and motif-based GRNs both performed similarly well on a benchmarking set consisting only of perturbations of TFs common to both data sources (Fig S3B). Combining the targets for the two GRNs led to an intermediate performance. However, several TF activity estimation methods allow stronger or more confident interactions in the GRN to be given extra weight. Giving extra weight to interactions present in both data sources did indeed improve the performance of the combined GRN over simply combining all targets (Fig S3B).

The ChIP-, motif- and eY1H-derived GRNs were integrated into a final network. Overall, this combined GRN covers 487 TFs, 58.5 % of the total estimated *C. elegans* TFs. This network also displayed improved performance when weighting interactions for appearance in multiple datasets (Fig 1E, Fig S3C). This combined and weighted GRN performs well, with an AUPRC of 0.722 and an AUROC of 0.641 (Fig 1E), a performance comparable to popular human GRNs like DoRothEA (MÜller-Dott *et al*. 2023). I call this final combined network *Cel*EsT (*<u>C</u>. <u>el</u>egans* <u>Es</u>timation of <u>T</u>F Activity; Table S3). The *Cel*EsT network performed similarly well on a benchmarking set without any correction for developmental age (Fig S4).

The ChIP, motif and eY1H datasets contributed similar numbers of TFs to *Cel*EsT (Fig 1F), although there were many more motif-derived interactions and few eY1H-derived interactions (Fig 1G). Although all data sources displayed good benchmarking performance in isolation, interactions were rarely shared (Fig 1G) even for those TFs that appeared in all three data sources (Fig S3D), in agreement with previous findings (Garcia-Alonso *et al*. 2019).

### Conservation of TF binding motifs across species identified high-confidence TF targets

While the integrated *Cel*EsT GRN described above performs well in the benchmarking pipeline, the high empirically-determined cut-offs for target inclusion led to a GRN with a large number of interactions; each TF has an average of 1053 targets (median 1397) and each target in turn is regulated by an average of 34.9 TFs (median 32). For researchers interested in identifying fewer, high-confidence TF targets, the benchmarking pipeline may offer an analytic resource to assess effective methods to distil these TF-target relationships into a more highly informative subset.

TF motif binding preferences are strongly conserved across even distantly related species, despite apparent divergence at the protein similarity level (Nitta *et al*. 2015). By comparison, individual motifs in promoters are vulnerable to even a single point mutation and thereby undergo evolutionary turnover in terms of loss and gain (TuĞrul *et al*. 2015). However, biologically important TF-promoter interactions will be subject to selection that leads to the retention (or reappearance) of a specific motif in orthologous target genes across species (Villar *et al*. 2014). As such, whereas strong matches to TF motifs within a promoter of a particular species may be spurious, the promoters of conserved TF targets are likely to feature the relevant motif across orthologous genes in multiple species more often than would occur by chance. Such a conservation-based approach to TF target identification has been previously employed in *C. elegans* (Glenwinkel *et al*. 2014).

Orthologous promoter sequences of *C. elegans* genes from 10 additional *Caenorhabditis* species were scanned for matches to known TF DNA-binding motifs (Fig 2A). I observed that 153/210 TFs present in at least 5 species have a significantly enriched overlap (Wilcoxon test, FDR < 0.2) of target orthologues with motif instances across species, consistent with the notion of conserved TF regulation. For each potential TF-target pair a conservation probability was calculated based on the random likelihood of identifying the observed pattern of motif conservation (both the number of species with significant motif matches and the strength of motif matches). Top motif matches for each TF in *C. elegans* were filtered by conservation probability (Fig S5A). TFs with few remaining targets after filtering (42/252 TFs) were removed. This led to a GRN which had a mean of 159.5 targets per TF (median 120.5). I observed that conserved motif-derived targets for those TFs with ChIP-seq data were 1.89 times more likely to feature a TF ChIP peak overlapping their promoter (chi-squared test, 95% CI 1.79-2.01, p-value 1.03 × 10^-103^), suggesting that they are indeed more likely to represent true TF targets.

**Fig 2.**
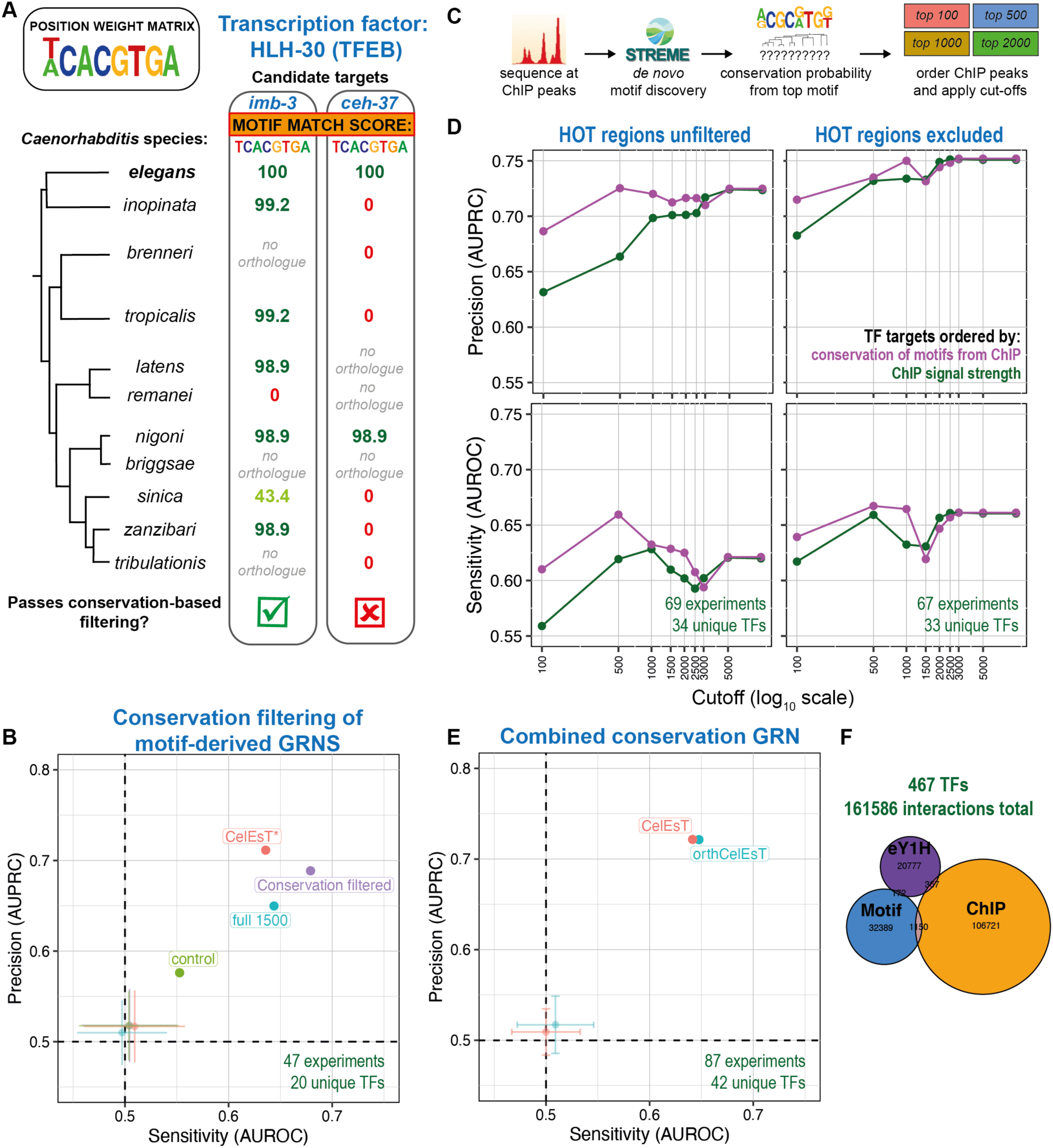
Identification of cross-species conservation of TF binding motifs in orthologous target genes improves network performance with fewer TF targets. **A)** Example of two potential target genes (*imb-3* and *ceh-37*) with a perfect match to the HLH-30/TFEB motif in their promoters in *C. elegans*. Looking across 10 additional *Caenorhabditis* species, the HLH-30 motif is found in the *imb-3* promoter in 6/7 species with one-to-one orthologues, whereas the motif is present in the *ceh-37* promoter in only 1/7 species. **B)** Benchmarking performance for motif-based GRNs after conservation-based filtering (purple) versus the best-performing *elegans*-only motif-based GRN (‘full 1500’, blue), a control network with the same number of targets per TF selected from the best *elegans* motif matches (‘control’, green) and the *Cel*EsT network subset to the same TFs (‘CelEsT*’, red). Axes show performance (AUROC/AUPRC) in benchmarking pipeline. Transparent points show mean AUROC/AUPRC of randomly shuffled networks; error bars show standard deviation. The number of experiments and unique TFs in the benchmarking set overlapping with TFs in the GRN is noted on the panel in green text. Results are shown for the best-performing multivariate model (‘mlm’) method; see also Fig S5B for performance with other methods. **C)** Schematic shows application of conservation-based filtering to ChIP-seq data. After a *de novo* motif search from sequences marked with ChIP-seq peaks, conservation probabilities are derived using the top resulting motif. Potential targets are then ordered by conservation probability, rather than ChIP peak signal strength as before, before applying cut-offs to compile GRNs. **D)** Line shows evolution of AUPRC (above) and AUROC (below) with increasing cut-off for maximum number of targets per TF for GRNs derived from ChIP-seq data with (right) or without (left) exclusion of target genes within HOT regions. Colours indicate network with targets ordered by conservation probability (purple) or ChIP peak signal strength (green). Note x-axis (cut-off) is on a log scale. Experiment/unique TF numbers in green text. See also Fig S5C. **E)** Benchmarking performance for orth*Cel*EsT, a combined GRN with conservation-based target prioritisation applied to regulons of both the motif dataset and the ChIP-seq dataset. Results are shown for the best-performing multivariate model (‘mlm’) method; see also Fig S5D for performance with other methods. **F)** Euler plot shows TF-target interactions derived from each dataset in the orth*Cel*EsT network (see also Fig S5E).

Despite featuring an order of magnitude fewer targets on average, this GRN performed better than the much larger motif-derived GRN using a cut-off of 1500 top *C. elegans* targets (Fig 2B, Fig S5B). Importantly, it very strongly outperformed a control GRN with a matched number of targets for each TF but consisting instead of the top motif matches in *C. elegans*. Thus, a cross-species conservation-based approach is very effective for filtering TF-target interactions to drastically reduce GRN size without loss of performance.

To test whether conservation-based target filtering could also be applied to ChIP data, *de novo* DNA-binding motifs were extracted from ChIP-seq data for 359 TFs (see Methods). Although conservation-based filtering of ChIP GRNs did not outperform unfiltered networks, it was observed that when TF targets were prioritised by conservation probability rather than by ChIP signal strength (Fig 2C), performance was substantially improved at lower cut-offs, whether HOT regions were excluded or not (Fig 2D). Indeed, for a ChIP-based GRN with HOT regions excluded, performance matched that of the unfiltered network at a regulon size cut- off of 1000 targets (Fig 2D; this cut-off affected 47 promiscuous TFs). Interestingly, GRNs compiled using *de novo* motifs derived from ChIP data performed markedly better than using the known DNA-binding motif where available (119 TFs; Fig S5C).

A combined GRN was created with both motif- and ChIP-derived targets filtered by conservation as described above, together with the eY1H dataset. This GRN performs almost identically to the *Cel*EsT network in the benchmarking pipeline (Fig 2E, Fig S5D), but with an average of 67% fewer targets per TF and a more equal balance between interactions derived from different datasets (Fig 2F, Fig S5E). However this comes at the loss of 20 TFs in the final network, for a total of 467. This combined conservation-based network is called orth*Cel*EsT (Table S4).

Though improving network performance, both exclusion of ChIP peaks in HOT regions and conservation-based filtering of motif occurrences led to loss of TFs from the final network. This may affect some researchers with a particular interest in a TF which was excluded but for which data nonetheless exists. With conservation-based ChIP target ordering affording greatly improved performance at low cut-offs without filtering out HOT regions (Fig 3D), it was possible to assemble a network with decent performance with no loss of TFs. Combining the top 500 conservation-prioritised ChIP targets together with the unfiltered top 1500 motif-derived targets and the eY1H data produces a network with 507 TFs in total. The performance of this network, called max*Cel*EsT (Table S5), compared favourably with *Cel*EsT, with a similar AUROC but somewhat lower AUPRC (Fig S5F).

**Fig 3.**
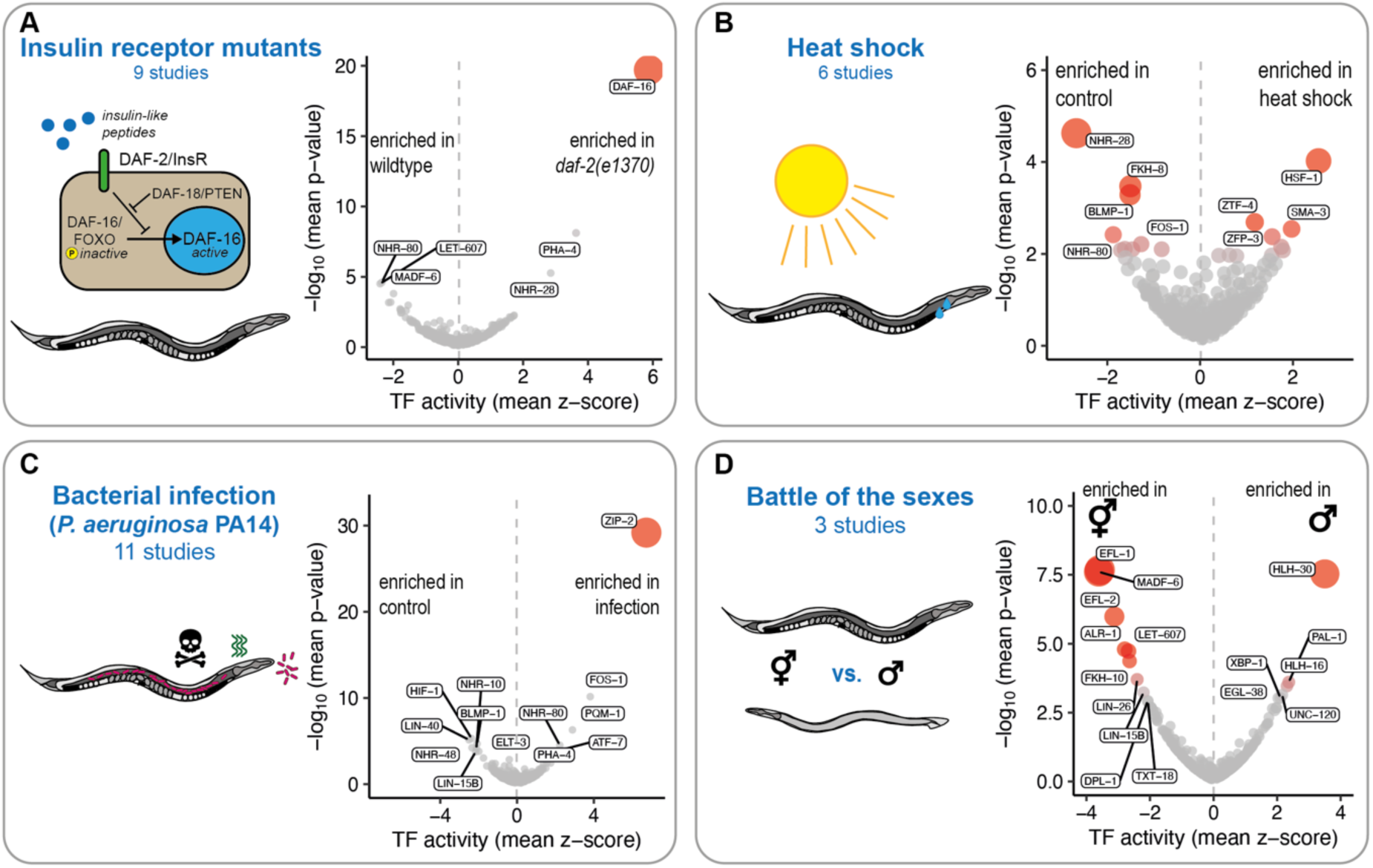
*Cel*EsT recapitulates known TF biology and generates new insights in commonly-studied genetic, environmental and physiological conditions. Volcano plots shows mean TF activity z-scores and geometric mean *p*-values across studies. The bubble size is proportional to the *p*-value. All analyses controlled for developmental age during differential expression analysis, with DE stats then analysed using the multivariate linear model TF activity estimation method and the *Cel*EsT network. **A)** Analysis of 9 studies comparing a severe mutant allele of the insulin receptor orthologue *daf-2* to wildtype controls. See also Fig S6, Fig S7A and Fig S8A. **B)** Analysis of 6 studies comparing heat-shocked animals to untreated control animals. See also Fig S7D and Fig S8B. **C)** Analysis of 11 studies comparing animals exposed to the pathogenic bacterium *Pseudomonas aeruginosa* PA14 to untreated controls. See also Fig S7E and Fig S8C. **D)** Analysis of 3 studies comparing the transcriptome of male animals to that of hermaphrodites. See also Fig S7F and Fig S8D.

### *Cel*EsT recapitulates known TF biology in intensively-studied conditions

To validate the performance of *Cel*EsT, TF activity estimation was performed on a variety of published RNA-seq datasets representing well-studied genetic, environmental and physiological conditions.

First, datasets from mutants of the insulin/IGF1-like signalling (IIS) pathway were analysed, focusing on the severe insulin receptor mutant *daf-2(e1370)* (Fig 3A, Fig S6). Amalgamating 9 studies (Fig S7A; Table S6), it was clear that activation (Fig 3A), though not upregulation (Fig S8A), of the FOXO orthologue DAF-16 was the major determinant of the transcriptional response to *daf-2* mutation, as is well established (Cahill *et al*. 2001; Lin *et al*. 2001). However a consistent and strong enrichment was also observed for differentially-expressed targets of PHA-4 and depletion of the targets of NHR-80 (Fig 3A). In 2 studies combining *daf-2* mutation with *daf-16* null mutations (Fig S7B) these changes are reversed, suggesting that these TFs act downstream of DAF-16 activity (Fig S6B). Although there is a significant DAF-16 ChIP peak in the promoters of both *pha-4* and *nhr-80*, neither gene is strongly regulated at the transcriptional level in *daf-2* mutants (Fig S8A). Similar reversed enrichments are found in data from mutation of the PTEN orthologue *daf-18* in a *daf-2* background (1 study, Fig S6C), which is known to supress *daf-2* mutant phenotypic defects (Ogg and Ruvkun 1998), and in data from *daf-16* mutants in a wildtype background (3 studies, Fig S7C, Fig S6D). The latter result implies significant baseline DAF-16 activity despite normal levels of IIS signalling.

Data was examined from 6 RNA-seq studies (Fig S7D; Table S6) of worms undergoing heat shock (Fig 3B). As expected, the strongest and most consistent enrichment was for HSF-1 (Heat Shock Factor 1; Fig 3B). *hsf-1* mRNA barely changed despite strongly increased HSF-1 activity (Fig S8B). More surprising was the even more significant depletion of the targets of NHR-28 (Fig 3B), suggesting that this nuclear hormone receptor has a major role in the transcriptional response to heat shock. Although *nhr-28* mutants have been shown to have strongly impaired survival of heat shock (Joshi *et al*. 2016), this role has been overlooked until now. As *Cel*EsT is an unsigned network, this depletion would also be consistent with an increased activity of NHR-28 in repressing target genes; however *nhr-28* itself appears to be downregulated at the transcriptional level upon heat shock (Fig S8B).

The transcriptional response to infection by pathogenic strains of the bacterium *Pseudomonas aeruginosa* (11 studies; Fig S7E) was strongly dominated by a single transcription factor, ZIP-2 (Fig 3C). ZIP-2 has been previously been described as one key mediator of the early transcriptional response to *P. aeruginosa* infection (Estes *et al*. 2010) among others including PQM-1 (Rajan *et al*. 2019) and ATF-7 (Fletcher *et al*. 2019). While these TFs also display increased activity, the extent to which ZIP-2 determines the transcriptional response to infection suggests that it is of primary importance. ZIP-2 and other TFs that have significantly increased activity exhibited strongly increased expression (Fig S8C), demonstrating that, unlike in heat shock, the response to *P. aeruginosa* infection is largely orchestrated by regulation of TFs at the transcriptional level.

Lastly, differences in TF activity between males and hermaphrodites/pseudo-females were examined (3 studies; Fig S7F, Table S6). The TF most strongly active in males relative to hermaphrodites was HLH-30 (Fig 3D), despite no difference in expression (Fig S8D). HLH-30 nuclear exit in hermaphrodites is induced by mating (Shi *et al*. 2019), which raises the possibility that this difference is not intrinsic but is instead due to low activity of HLH-30 in hermaphrodites raised in the presence of males. I also found strongly increased activity of PAL-1 (Fig 3D), which is known to be important for tail development in males, despite a lower *pal-1* expression level (Fig S8D). In hermaphrodites the E2F-like TFs EFL-1 and EFL-2 and their putative cofactor DPL-1 were more active (Fig 3D), likely due to crucial roles in the hermaphrodite germline (Chi and Reinke 2009).

In sum, these results both validate the use of *Cel*EsT for TF activity estimation and highlight its potential to generate novel insights.

## DISCUSSION

TF activity estimation from target gene expression is a potent and increasingly popular method that allows inference of mechanistic insights from transcriptomic data. However without prior knowledge of TF targets this method cannot be applied. Here, inspired by the human GRN DoRothEA (Garcia-Alonso *et al*. 2019), disparate large-scale experimental datasets previously generated by *C. elegans* researchers were integrated to provide a unified knowledge resource, the *Cel*EsT gene regulatory network, thus making this powerful analysis available to the worm community. Importantly, *Cel*EsT has been benchmarked against a comprehensive set of TF target perturbation RNA-seq experiments to ensure optimal processing and integration of distinct datasets.

*CelEsT* can be used with the *decoupleR*/*decoupler* package (Badia-i-Mompel *et al*. 2022) in either *R* or *Python* to infer TF activity estimates from gene expression data within minutes on a standard computer. I provide an annotated and customisable *R* script to users who wish to use *Cel*EsT with specific parameter settings or alternative methods (File S1 and S2 respectively). However an *R Shiny* application is also provided at github.com/IBMB-MFP/CelEsT-app so that members of the community can perform analysis with minimal coding experience required. The *Cel*EsT app permits input of either existing differential expression (DE) statistics or raw sample gene-level read counts. In the latter case, the *Cel*EsT app can perform differential expression analysis and also allows the user to opt to correct for developmental age in order to eliminate any spurious differential gene expression derived from effects of their treatment on development (Bulteau and Francesconi 2022).

Although multiple methods of TF activity estimation were tested, it was shown that for these networks the multivariate linear model (‘mlm’) method consistently performed best. Importantly this method considers multiple TFs in a single calculation, which allows for disentangling distinct effects of multiple TFs even where their targets exhibit significant overlap. This method also happens to be the fastest and least computationally demanding of those tested. As such, it is recommended that users that employ the GRNs provided here use the ‘mlm’ method of TF activity estimation for maximum performance.

The principal aim of this study was not to provide exhaustive lists of targets for each TF, but rather to find a core functional set of targets for each TF that provided for accurate estimation of its activity. While the use of the *Cel*EsT GRN is recommended, the orth*Cel*EsT and max*Cel*EsT networks are also presented, which minimise TF-target interactions and TF loss at the cost of TF numbers and performance, respectively. The orth*Cel*EsT GRN, with approximately 1/3 of the interactions of the main *Cel*EsT network, represents a valuable resource for those seeking high-confidence TF targets in *C. elegans*. Meanwhile, the max*Cel*EsT GRN may be useful to researchers with an interest in a particular TF which was ultimately excluded from the *Cel*EsT network.

Assigning targets to TFs based on putative matches to DNA-binding motifs is inherently likely to produce false positives (Garcia-Alonso *et al*. 2019). Here it was shown that using conservation-based target filtering in combination with TF activity estimation benchmarking is a powerful method to winnow down potential motif-based targets to a highly informative set for each TF. This is likely effective precisely because it helps to eliminate false positives.

Conservation of TF-binding motifs across species in the promoters of their targets is a strong indication that a given target does not represent a spurious sequence match but is a biologically-relevant target that leads to selectable phenotypic impacts when TF regulation is lost (TuĞrul *et al*. 2015). Importantly, this method is highly applicable to a host of model organisms; in the age of comparative genomics it is common that multiple congeneric species of favourite models have available genome sequences (HernÁndez-Plaza *et al*. 2023).

It was also shown that a similar approach is applicable to ChIP-seq data to highlight biologically-relevant targets based on conservation of *de novo* motifs. Targets marked by ChIP peaks are less likely than those with putative motif instances to be false positives, and strong filtering of ChIP-based targets according to *de novo* motif conservation did not improve network performance. However motif conservation was a more effective metric than ChIP peak signal for target prioritisation when applying stringent cut-offs to the number of targets per TF. Surprisingly this method was effective despite using only one of several *de novo* motifs, which often did not correspond to *in vitro*-determined motifs where known (see Methods). The top *de novo* motifs discovered may not represent the direct sequence preference of a TF but likely also include motifs for relevant regulatory co-factors (Liu *et al*. 2018). Regardless, the higher performance observed using *de novo* motifs over known motifs where applicable shows that *de novo* motifs nonetheless encode a potent biological signal which aids in identifying the most important targets of a TF.

One important limitation of *Cel*EsT is that it is an unsigned network that does not assign a mode of regulation (i.e. activation, repression or bifunctional) to each TF. As such, greater activity of a repressor will lead to a depletion of its target mRNA and thus to apparently lower estimated activity. Further, TF estimation activity methods are known to be less effective for bifunctional TFs (Garcia-Alonso *et al*. 2019). While a subset of *C. elegans* TFs have an assigned mode-of-regulation in WormBase or UniProt (Table S1), this annotation is likely to be far from complete and its inclusion somewhat impaired benchmarking performance (not shown). In any case, the tendency towards positive correlations between TF expression and activity (Fig S8) suggests that the dominant trend is towards positive TF regulation of targets.

Although comprehensively representing the bulk of *C. elegans* TF perturbation RNA-seq experiments conducted to date, another weakness is the limited size of the benchmarking set, both in terms of experiments and unique TFs represented. It is likely that with a larger benchmarking dataset and greater TF coverage, benchmarking performance would be less noisy and more likely to result in network parameters that provide optimal performance across all TFs. Expansion of a benchmarking dataset by future efforts towards a consortium or database providing systematic transcriptomic characterisation of TF knock-out/knock-down animals would pay dividends for the *C. elegans* community.

In conclusion, the introduction of *Cel*EsT to enable TF activity estimation is an invaluable addition to the toolbox of the worm community. In addition, this study provides a roadmap for researchers who wish to effectively utilise existing knowledge resources for TF activity estimation in other model organisms. I hope that *Cel*EsT will not only enable more effective interpretation of future experiments but also allow a re-evaluation of past experiments to generate many novel insights, even from old data.

## Acknowledgments

This work was funded by a Ramon y Cajal fellowship (RYC2021-034496-I) awarded by Spain’s Ministerio de Economía y Competividad . I thank Alexandra Avgustinova, Andre Faure, Mirko Francesconi and Adam Klosin for critical comments on the manuscript. Thanks to Charlie Cotton for inspiring a suitably celestial logo, to my niece Celeste Perez for inspiring the name, to Jennifer Semple for use of her excellent worm illustrations and to Pau Badia-i-Mompel for always helpful, polite and rapid responses to questions posted on the *decoupler* GitHub issues page. Fig S2D contains the Protein of the Month illustration of HIF-1 (credit: David S. Goodsell and the RCSB PDB) modified in agreement with the conditions of its <u>CC-BY-4.0 license</u>.

## Data and code availability

No original datasets were produced in this study. Availability of all datasets used is described in Methods and Tables S2, S6 and S7. All analysis code is present on Github at github.com/IBMB-MFP/CelEsT-MS.

## METHODS

### Assembly of TF perturbation benchmarking RNA-seq dataset

To produce a set of TF perturbation RNA-seq experiments for benchmarking, I used NCBI’s *eUtils* tool suite to download summaries of all *C. elegans* RNA-seq experiments in the Gene Expression Omnibus (GEO) accessed with the following query:

> "Caenorhabditis elegans"[Organism] AND "expression profiling by high throughput sequencing"[DataSet Type].

The search was conducted on the 15/01/2024 and yielded 882 entries. I examined each entry for experiments where the expression of a transcription factor present in any of the three principal datasets was perturbed (either knockdown or overexpression) and that had matched control samples. A few intensively studied TFs were over-represented (e.g. DAF-16); to ensure that the benchmarking set was not too heavily skewed towards a small number of TFs I arbitrarily limited the number of individual experiments for any single TF to a maximum of 4, choosing to retain experiments that represented high quality datasets (in terms of replicates and sequencing depth) and a maximal diversity of perturbations, conditions and genetic backgrounds. The result was a set of 90 experiments for 45 unique TFs from 73 distinct studies (Table S2). Note that a few of these TFs were ultimately ‘lost’ from the tested GRNs due to having very few targets remaining after e.g. HOT region exclusion, such that the maximum size of benchmarking set actually used in the study was 87 experiments for 42 unique TFs.

For each study I noted the identifiers for TF-perturbed and control samples and downloaded the raw sequencing data using NCBI’s command-line package *SRA toolkit*. Reads were aligned with the *bowtie2* (Langmead and Salzberg 2012) package to the *C. elegans* genome (version WS288). Gene-level read counts were computed with “featureCounts” function (Liao *et al*. 2014) from the *Subread* package by cross-referencing to the WS288 canonical geneset ‘gtf’ file downloaded from WormBase. Gene-level read counts were imported into *R* and differential expression analysis was performed with the *DESeq2* package(Love *et al*. 2014).

I controlled for potential mismatches of developmental speed between control and treatment samples using the *RAPToR* package (Bulteau and Francesconi 2022), following the procedure outlined in the *RAPToR* vignette. Briefly, sample ages were estimated using *RAPToR* against the references provided in the associated *wormRef* package; appropriate references were selected according to the annotated developmental stage on GEO or the relevant paper. I then interpolated counts from the appropriate reference for a range covering one hour before and after the age range estimated to be covered by the test samples. I estimated gene dispersions using *DESeq2* using only the test samples and then created a *DESeq2* object combining these dispersions with the test counts and interpolated counts from the reference. I then performed the DE analysis by fitting a model including the effect of time modelled by a spline, treatment (TF perturbation vs control) and batch (test samples or interpolated reads). The DE statistics for contrasting TF perturbation vs control samples were extracted – these statistics exclude any differential expression due to developmental differences alone. Note DE analysis for two experiments on dauer larvae were conducted without accounting for developmental difference due to the lack of an appropriate reference on which to stage the samples. The DE statistics for all genes (not only those significantly differentially expressed) within each experiment constituted the benchmark dataset used in the benchmarking pipeline. Prior to use in the benchmarking pipeline I reversed the sign of DE statistics from experiments involving TF overexpression, such that the directions of all perturbations appeared equal.

### Additional data resources excluded from consideration

An additional potential source of information about TF-target interactions was curated user-supplied regulatory interactions from the worm community platform WormBase (Harris *et al*. 2020), but these interactions are very few relative to those supplied by other resources and often represent indirect relationships. As inclusion of these interactions did not improve the performance of GRNs in the TF activity estimation benchmarking pipeline (data not shown), I did not consider the WormBase interactions any further. I also attempted to infer regulatory relationships by TF-target co-expression using SJARACNe (Khatamian *et al*. 2019), using as input >600 *C. elegans* wild isolate RNA-seq samples (Zhang *et al*. 2022), but I found the resulting GRN had no predictive capacity (Fig S9). Likewise, I extracted TF mode of regulation (i.e. whether activator or repressor) where available from gene descriptions in WormBase and UniProt. However this information did not consistently improve network performance and so was discarded. As such the resulting networks from this study are unsigned i.e. they do not assign a mode of regulation to each TF. Nonetheless the TF modes of regulation compiled from WormBase and UniProt can be found in Table S1.

### Assembly of GRNs from TF ChIP data

I downloaded the processed ChIP-seq TF-binding data for all experiments reported by the modERN consortium (Kudron *et al*. 2018; Kudron *et al*. 2024) from epic.gs.washington.edu/modERNresource/. 6 TFs are listed in the Supplementary Materials of (Kudron *et al*. 2024) as having been successfully ChIPed but were not present in the Peaks file, presumably due to erroneous omission. Of these, 4 were present on the ENCODE platform (B0019.2, W05B10.2/CCCH-3, T26A8.4, W05H7.4/ZFP-3); I downloaded the processed narrowPeak files from the ENCODE website. 2 additional TFs (W03F9.2, Y116A8C.19) were present in the original modERN release (Kudron *et al*. 2018) (epic.gs.washington.edu/modERN) but were absent from the latest release, but without any sign that the data had been revoked or that they were no longer considered TFs. As such, they were included in this study and the data was obtained from the original modERN release.

Additionally I downloaded summaries of all *C. elegans* ChIP-seq experiments from GEO (accessed 04/02/2024), a total of 604 experiments. I examined each entry for pulldowns of TFs and matched input controls. Of note, a number of early experiments from modENCODE/modERN were found to be present in GEO but these were ignored. In some cases the data was already present in the modERN resource platform; in some cases it was not present but the ChIP for the TF was annotated as failed in the Supplementary Material of (K^UDRON^ *et al*. 2024) or marked as revoked on the ENCODE platform. I identified ChIP data for an additional 6 TFs (F22A3.5/CEH-60, F56F11.3/KLF-1, T05C1.4/CAMT-1, T23D8.8/CFI-1, ZC376.7/ATFS-1 and Y51H4A.17/STA-1); however the data for CFI-1 was ultimately excluded due to an unusually high number of binding targets. Raw sequencing data for TF ChIP and input controls were downloaded with *SRA toolkit* (E^DWARDS^ 2021) and aligned to the *C. elegans* genome (WS288) with *bowtie2*. I then used the *Genrich* package to call peaks, as *Genrich* allows integrated peak-calling against a control file using multiple replicates. The resulting narrowPeak files were then treated as those downloaded from ENCODE.

Potential targets of TF ChIP peaks were assigned manually, ignoring the annotated targets present in some processed files from modERN. From the *TxDb.Celegans.UCSC.ce11.refGene* package/object in *R* (T^EAM^ 2019) I obtained genomic locations for protein-coding genes. As the objective was to have potential targets with robust and differentiated expression in order to be useful as markers for inferring TF activity, I censored these potential genes by those whose expression was detected in at least 2/3 of the ∼600 samples in the largest available *C. elegans* RNA-seq dataset (triplicate samples from ∼200 wild strains of the *C. elegans* Natural Diversity resource (CeNDR); (Z^HANG^ *et al*. 2022). I also eliminated those genes that were annotated as being downstream in a C. elegans operon in {Allen, 2011 #48), as they would not likely be controlled by proximal promoter regions. This resulted in a set of 14083 potential target genes. I derived the genomic regions taken as corresponding to proximal promoter regions for these genes by taking 1000 bp upstream and 200bp downstream of the transcription start site (TSS) using the “genes()” and “promoters()” functions of the *GenomicFeatures* package (L^AWRENCE^ *et al*. 2013) in *R*. For each ChIP experiment I then used the “findOverlaps()” function of the *GenomicRanges* package (L^AWRENCE^ *et al*. 2013) in *R* to identify overlaps between ChIP peaks and these candidate target promoter regions. Of note, multiple interactions could derive from a single ChIP peak if it fell in a region where two promoters overlapped; i.e. rather than assigning each individual peak to a single definitive target as in the processed modERN files, I counted all TFs bound to each promoter as potential regulators regardless of any greater proximity of a peak to the TSS of a different gene.

ChIP peaks for a given TF were prioritised into an ordered list based first on repeated occurrence across multiple independent experiments, if applicable, and then based on the strength of the ChIP peak obtained from the narrowPeak files. Cut-offs were then applied, taking the top targets corresponding to a given cut-off from this ordered list, to generate distinct GRNs for benchmarking. Any TFs with fewer than 15 targets were filtered out of the final GRN.

Genomic locations of High Occupancy Target (HOT) regions were taken as defined in Table S10 of (K^UDRON^ *et al*. 2018). Of note, this table listed the number of experiments for which a given region was bound by TF but repeated occurrence of binding of the same TF in multiple experiments were considered as independent instances counting towards this total and thus to the definition of a HOT region. As I did not consider that a region should be excluded in part due to consistent binding of those TFs which had multiple experiments, I instead counted the number of distinct TFs, of the total of 217 assayed, annotated as binding the HOT region in any experiment. GRNs were generated excluding as potential targets any gene whose promoter overlapped a HOT region, defined by binding of a given cut-off of distinct TFs out of the total 217 assayed in that study. As shown in Fig S1D, the best performance (considering trade-offs with ‘loss’ of those TFs with few remaining targets from the network) came from applying an exclusion criterion of 50/217 TFs to regions considered to be HOT regions.

### Assembly of GRNs from *in vitro* DNA-binding motifs

I downloaded all available DNA-binding motifs for *C. elegans* TFs from CisBP (Weirauch *et al*. 2014) (accessed 23/02/2023). I excluded any motifs derived from ChIP-seq data, retaining only those obtained via *in vitro* methods (principally protein-binding microarrays (PBMs) and SELEX experiments). Some motifs were directly measured from that TF’s DNA-binding domain, while others were indirectly inferred from similar DNA-binding domains from other species. I excluded any indirect motifs that derived from directly-measured motifs of *C. elegans* TFs to avoid motif duplication. Likewise, in the case where a single motif from another species was assigned to multiple *C. elegans* TFs, I manually selected one TF based on criteria of best similarity score in CisBP and expression level in *C. elegans* and excluded the other TFs.

Candidate target promoter regions were identified as described above. The sequences corresponding to these promoters were obtained from the *BSgenome.Celegans.UCSC.ce11* package/object using *GenomicRange’s* “get_sequence()” function in *R*. I ran the FIMO tool from the *MEME Suite* collection of sequence motif tools (Bailey *et al*. 2015), searching for instances of the CisBP TF motifs in the promoter sequences. Promoters were ordered according to the best-scoring motif match returned by FIMO before cut-offs were applied to generate GRNs. For calculating homotypic binding scores taking into account additional matches beyond the first, points for additional matches were added to the score for the best match with exponentially diminishing returns for additional matches. The score of the second-best match was added divided by 5^1^, of the third-best match divided by 5^2^ and so on. The diminishing returns parameter 5 was selected as the best-performing by benchmarking GRNs with scores computed with different diminishing returns parameters (not shown).

### Assembly of GRNs from eY1H data

eY1H data was taken from Table S2 of (Fuxman Bass *et al*. 2016). I filtered interactions to only those annotated as belonging to the ‘high-quality dataset’. This gave 21864 interactions for 366 unique TFs. After filtering out TFs with fewer than 15 targets, there remained 20773 interactions for 160 TFs.

### Assembly of GRNs from SJARACNe-inferred co-expressed TF/gene pairs

Sample ages for ∼600 RNA-seq samples from CeNDR wild *C. elegans* strains (Zhang *et al*. 2022 were inferred using RAPToR from TPM-normalised values downloaded as a supplementary file from GEO entry GSE186719. Sample raw transcript-level counts (also obtained as a supplementary from the same GEO entry) were summed to generate gene-level counts, which were then processed with a variance-stabilising transformation using the “vst()” function of DESeq2 in R. The vst-normalised values were modelled as a function of time with the “smooth.spline()” function in *R* with 6 degrees of freedom to remove any differences between samples due to differential developmental age. The residuals were extracted from the model and used as input values for SJARACNe {Khatamian, 2019 #21). Note I also repeated this analysis with values with no correction for developmental age without any improvement in performance (not shown). I filtered all genes to remove any genes that were not detected in every sample. I supplied a list of nodes (TFs) to SJARACNe that consisted of all of the TFs present in any of the other three datasets, except those not detected in every sample (leading to exclusion of 123 TFs). I ran SJARACNe on the command line and filtered the output for interactions with a p-value below 0.0001. A GRN was then constructed with the remaining interactions for each TFs, weighted according to the sign of the interaction detected by SJARACNe.

### Assembly of combined GRNs from multiple data sources

When combining GRNs from different datasets, I always used unfiltered GRNs (i.e. that did not have TFs excluded due to low target numbers); this was because a TF might be able to reach the threshold for inclusion (15 targets) by combining targets from multiple data sources, although it had failed to reach this threshold so in any single data source. For unweighted GRNs target lists for each TF were simply combined (with removal of duplicate interactions); for weighted GRNs interactions were given weights according to their appearance in the three datasets. Interactions appearing in all three data sources were given a maximum weight of 1, while interactions appearing in 2 or 1 data sources were given weights of 0.67 and 0.33 respectively.

### Benchmarking

In order to benchmark the performance of the GRNs against our TF perturbation benchmarking set, I used the *decoupler* package implemented in *Python* (Badia-i-Mompel *et al*. 2022). Note that the benchmarking pipeline is more fully developed and much faster in the *Python* version of the *decoupler* package relative to the *R* version. I used the “benchmark()” function, using the methods ‘mlm’, ‘wlm’, and ‘wsum’, also computing a multi-method consensus score. In order to compare the benchmarking output against randomly shuffled networks, I used the “shuffle_net()” function of *decoupler* to produce 100 randomly shuffled networks for each of our tested GRNs. Benchmarking was then repeated for each of the shuffled networks. The plotted points and error bars for random networks represent the mean and standard deviation of the performance metrics (AUROC/AUPRC) for the 100 shuffled iterations of that network.

### Cross-species conservation-based filtering of TF targets

In order to perform cross-species conservation-based filtering of potential TF targets by I first build *BSGenome* (Pagès 2024) and *TxDb* packages in *R* for each of the 10 additional *Caenorhabditis* species (*brenneri, briggsae*, *inopinata*, *latens*, *nigoni*, *remanei*, *sinica*, *tribulationis*, *tropicalis* and *zanzibari;* genome versions in Table S7) in order to obtain sequences corresponding to potential target promoters. The genome assemblies for these species were downloaded as a FASTA file from WormBase ParaSite (Howe *et al*. 2017). Chromosome/contig numbers were extracted and the genome-sequence FASTA file for each species was split into individual FASTA files for each chromosome/contig. I then created a seed file and used *BSGenome*’s “forgeBSgenomeDataPkg()” function to create a ‘BSGenome’ data package for each species. I also created a ‘TxDb’ genome annotation object for each species using annotation files downloaded from WormBase ParaSite using *GenomicFeatures*’ “makeTxDbFromGFF()” function.

I then obtained the genomic locations of the promoters of genes as described above for *C. elegans* using the “promoters()” and “genes()” function of *GenomicFeatures* with the corresponding species’ ‘TxDb’ annotation data object. I filtered out any promoters which were incomplete due to proximity to the edge of a chromosome/contig. For each species I restricted the set of potential target promoters to those that were annotated as one-to-one orthologues of a *C. elegans* gene in Ensembl *BioMart* (Kinsella *et al*. 2011) queried using the *biomaRt* package in *R*. I also filtered out candidate promoters that were annotated as downstream in an operon in either *C. elegans* as described above or *C. briggsae* (annotations for the latter obtained from Table S7 of (Jhaveri *et al*. 2022)), as operons show a considerable degree of conservation across *Caenorhabditis* species (Qian and Zhang 2008; Pettitt *et al*. 2014) and so genes contained downstream within operons in at least one *Caenorhabditis* species are less likely to be controlled by the sequences proximal to the annotated TSS. I then obtained the sequences for the set of filtered promoters using “get_sequences()” and the corresponding species ‘BSGenome’ data object.

I then searched for all *C. elegans* TF DNA-binding motifs obtained from CisBP as above on the promoter sequences for each species using FIMO as described above. For each TF I extracted the best motif score for each potential target promoter in each species. If the potential target promoter sequence was absent in a species (due to the absence of an annotated one-to-one orthologue or due to filtering out incomplete promoters) then it was marked with NA in that species; if a potential target was present but exhibited no motif match it was marked with a 0. If there was not a one-to-one orthologue of the TF itself in that species, then all potential targets were marked with NA for that TF in that species. I calculated a conservation score as follows. First, I transformed the best motif match score from FIMO for each target into a rank-quantile for that motif and that species (the promoter with the best matching motif score in a genome had a rank-quantile close to 0, whereas promoters with no motif match had a rank-quantile of 1). I then computed the product of these rank-quantiles for each TF-promoter combination across all species to obtain the conservation score for that target and that TF.

I calculated the probability of conservation of a motif’s presence across species in orthologous targets as follows. For each TF, the rank-quantiles of promoters in each species were shuffled randomly between all non-NA targets (thus preserving the distribution of motif scores within each species and the number of species considered for each promoter) . An empirical random distribution of conservation scores was generated by calculating the product of the rank-quantiles for 10000 iterations of random shuffling. The observed conservation scores were compared to this random distribution to obtain an empirical p-value for each TF-target pair. These empirical p-values were adjusted for multiple comparisons using the Benjamini Hochberg/ method to generate FDR values.

I generated multiple GRNs using different initial cut-offs of *C. elegans* scores and then filtering down to those interactions with orthology-derived conservation probabilities meeting different FDR cut-offs. These GRNs were benchmarked to find the optimal cut-offs for *C. elegans* score and conservation probability. As the conservation-based GRN with the best performance used a seemingly high FDR of 0.5, these FDRs are likely conservative.

### Ordering of ChIP targets by *de novo* motif conservation across orthologues

I applied the “STREME” algorithm (BAILEY 2021) from *MEME Suite* by running the function “runStreme()” from the *R* package *memes* (Nystrom and Mckay 2021). As input sequences I used the 100 bp surrounding the centre of ChIP peaks, which was calculated by adding the ‘peak’ value (10^th^ column of ENCODE narrowPeak files) to the ‘start’ genomic co-ordinate for each peak. A minimum of 100 ChIP peaks were used to extract *de novo* motifs.

To judge the performance of STREME in defining *de novo* motifs, I used the “Tomtom” algorithm from *MEME Suite* to compare *de novo* motifs to directly-determined motifs from CisBP. I observed much better performance when excluding ChIP peaks located in HOT regions (cut-off of 50 applied as described above), as found elsewhere (data not shown; (Gerstein *et al*. 2010; Kudron *et al*. 2024)). I also experimented with using a maximum of the top 300 ChIP peaks, rather than all peaks, for the *de novo* motif search with STREME. Restricting the ChIP peaks in this way led to fewer motifs discovered per TF (mean of 4.32 vs 8.08 with all peaks). Although the motif discovery rate (defined as the identification of at least one motif with a significant similarity to the known motif) was lower (27/77 TFs vs 31/77), the known motif was the top hit more often (14/77 TFs vs 12/77 TFs). As such I restricted the ChIP peaks for *de novo* motif discovery to the top 300 non-HOT peaks.

I proceeded to calculate conservation probabilities as described above using the top *de novo* motif for each TF. Despite the top *de novo* motif matching the known *in vitro* motif less than 20% of the time, ChIP GRNs with targets ordered according to conservation probability performed surprisingly well.

Additionally I ordered ChIP targets by the conservation probabilities for the known motif, rather than the *de novo* motif, for the 119 TFs present in both the ChIP data and CisBP (either directly- or indirectly-determined motifs). Interestingly these networks performed markedly worse than those using the top *de novo* motif.

### Application of *Cel*EsT to RNA-seq datasets for genetic, environmental or physiological conditions

I identified candidate datasets for insulin signalling mutants, heat shock, bacterial infection with pathogenic *P. aeruginosa* strain PA14 and males vs hermaphrodites from the *C. elegans* RNA-seq datasets in the GEO. RNA-seq datasets were downloaded, aligned and gene-level read counts computed as described for the preparation of the benchmark dataset. Similarly, DE analysis accounting for developmental age was conducted as previously described. TF activity estimates were derived for each experiment using *Cel*EsT with *decoupleR* in *R* with the experiment’s DE stats as input and using the ‘mlm’ method. I plotted TF activities as heatmaps and generated study correlation matrices of TF activities to detect and exclude any outlying samples (*daf-2(e1370)* mutants one study – GSE36041; *daf-16* null mutants two studies – GSE240821 & GSE108848; heat-shock one study – GSE122015; males vs hermaphrodites one study – GSE222447).

To amalgamate the results of the independent experiments for each condition, I converted the TF activities for each experiment into a z-score by subtracting from each TF activity score the mean of the scores for all TFs and dividing by the standard deviation of the scores. I then took the mean activity z-score for each TF across all experiments for a condition. Similarly, for each TF I took the geometric mean of the p-values computed in each experiment. Volcano plots show these mean activity z-scores against the -log_10_(geometric mean p-value).

### Plotting

Plots were rendered in *R* using the *ggplot2* (line/volcano/scatter plots), *gplots* (heatmaps) and *eulerr* (Euler plots) packages.

---

**Fig S1.**
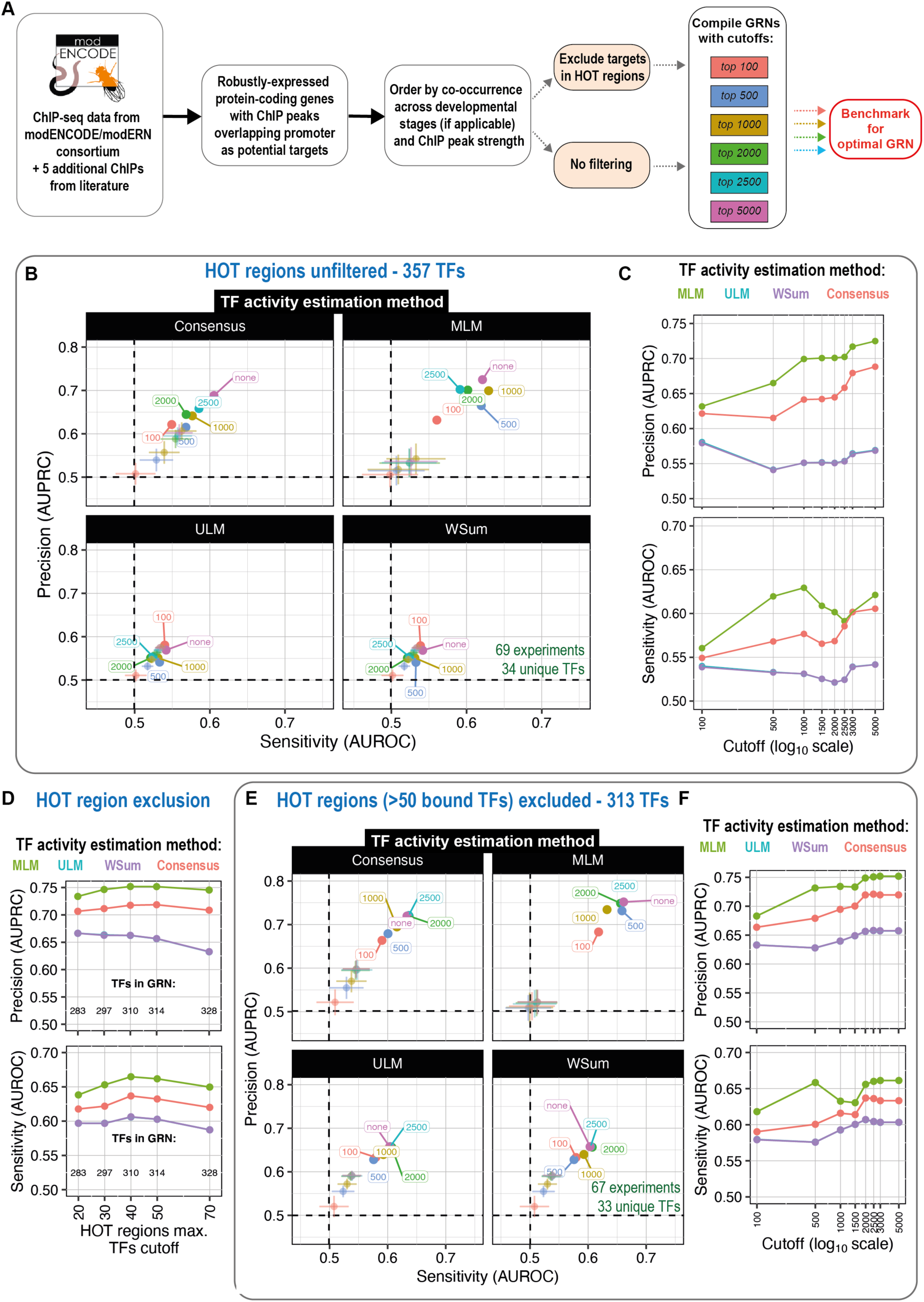
ChIP-derived GRNs perform well when potential targets in HOT regions are excluded (related to Fig 1) **A)** Schematic of pipeline to generate GRNs from ChIP-seq datasets. **B)** Benchmarking performance with different TF activity estimation methods (subpanels) for GRNs derived from ChIP-seq data with no filtering of High Occupancy Target (HOT) regions. Axes show performance (AUROC/AUPRC) in benchmarking pipeline. Colours show different cut-offs for maximum number of targets per TF. ;Colours correspond to cut-offs; transparent points show mean AUROC/AUPRC of randomly shuffled networks; error bars show standard deviation. The number of experiments and unique TFs in the benchmarking set overlapping with TFs in the GRN is noted on the panel in green text. **C)** Line shows evolution of AUPRC (above) and AUROC (below) with increasing cut-off for maximum number of targets per TF for GRNs derived from ChIP-seq data with no filtering of HOT regions. Colours indicate TF activity estimation methods. Note x-axis (cut-off) is on a log scale. Fig 1B shows benchmarking performance for AUPRC and ‘mlm’ only. **D)** Line shows evolution of AUPRC (above) and AUROC (below) with increasing cut-off for maximum number of distinct TFs (out of 217 from (Kudron *et al*. 2018)) bound to a potential HOT region for overlapping target genes to be retained. Colours indicate TF activity estimation methods as in panel C. More stringent (i.e. lower) cut-offs lead to exclusion of more TFs from the GRN that retain fewer than 15 targets outside of these regions; text shows the number of TFs retained for each cut-off. No cut-off for maximum number of targets per TF was applied for these GRNs. Benchmarking set consisted of 66 / 32 (for HOT region cut-off ≤ 30) or 67 / 33 (cut-off > 30) experiments and TFs respectively. **E)** Benchmarking performance with different TF activity estimation methods for GRNs derived from ChIP-seq data with target promoters overlapping HOT regions excluded. Details as for panel B. **F)** Evolution of AUPRC (above) and AUROC (below) with increasing cut-off for ChIP-derived GRNs with HOT regions excluded for all methods. Fig 1B shows benchmarking performance for AUPRC and ‘mlm’ only. mlm: multivariate linear model, ulm: univariate linear model, wsum: weighted sum

**Fig S2.**
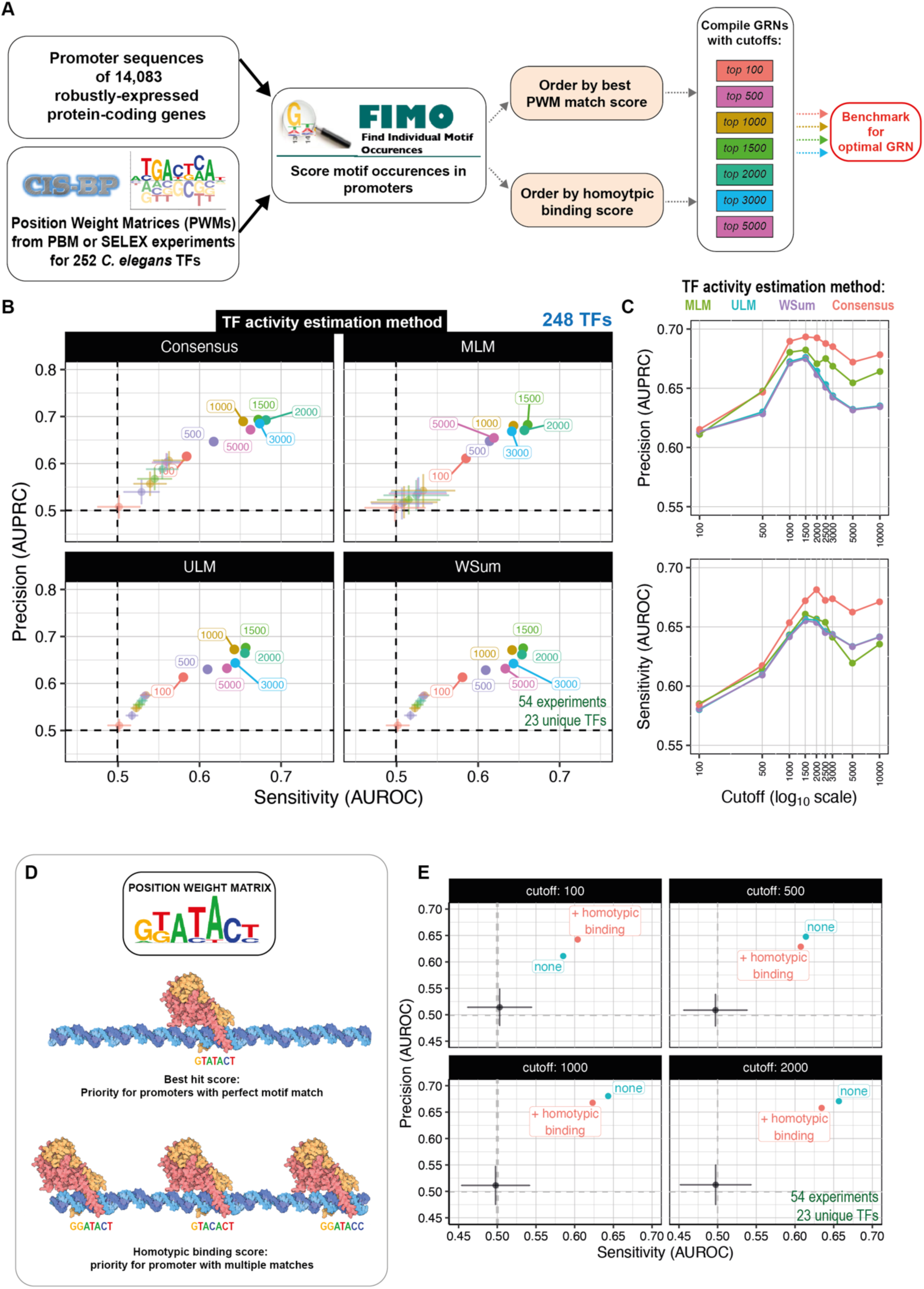
GRNs derived from promoters with best motif matches perform well with an optimal cut-off of regulon size (related to Fig 1) **A)** Schematic of pipeline to generate GRNs from PBM/SELEX-derived TF DNA binding motif datasets. **B)** Benchmarking performance with different TF activity estimation methods (subpanels) for GRNs derived from DNA binding motifs. Axes show performance (AUROC/AUPRC) in benchmarking pipeline. Colours show different cut-offs for maximum number of targets per TF. ;Colours correspond to cut-offs; transparent points show mean AUROC/AUPRC of randomly shuffled networks; error bars show standard deviation. The number of experiments and unique TFs in the benchmarking set overlapping with TFs in the GRN is noted on the panel in green text. Fig 1C shows benchmarking performance for AUPRC and ‘mlm’ only. **C)** Line shows evolution of AUPRC (above) and AUROC (below) with increasing cut-off for maximum number of targets per TF for GRNs derived from derived from DNA binding motif. Note x-axis (cut-off) is on a log scale. **D)** Schematic illustrates standard pipeline (considering only the best motif match per promoter) or homotypic binding cluster pipeline (adding extra priority to promoters with multiple matches which may be imperfect). **E)** Benchmarking performance for different cut-offs (subpanels) with targets ordered by best match or by a homotypic binding score which takes into account multiple matches per promoter. Details as for panel B. mlm: multivariate linear model, ulm: univariate linear model, wsum: weighted sum

**Fig S3.**
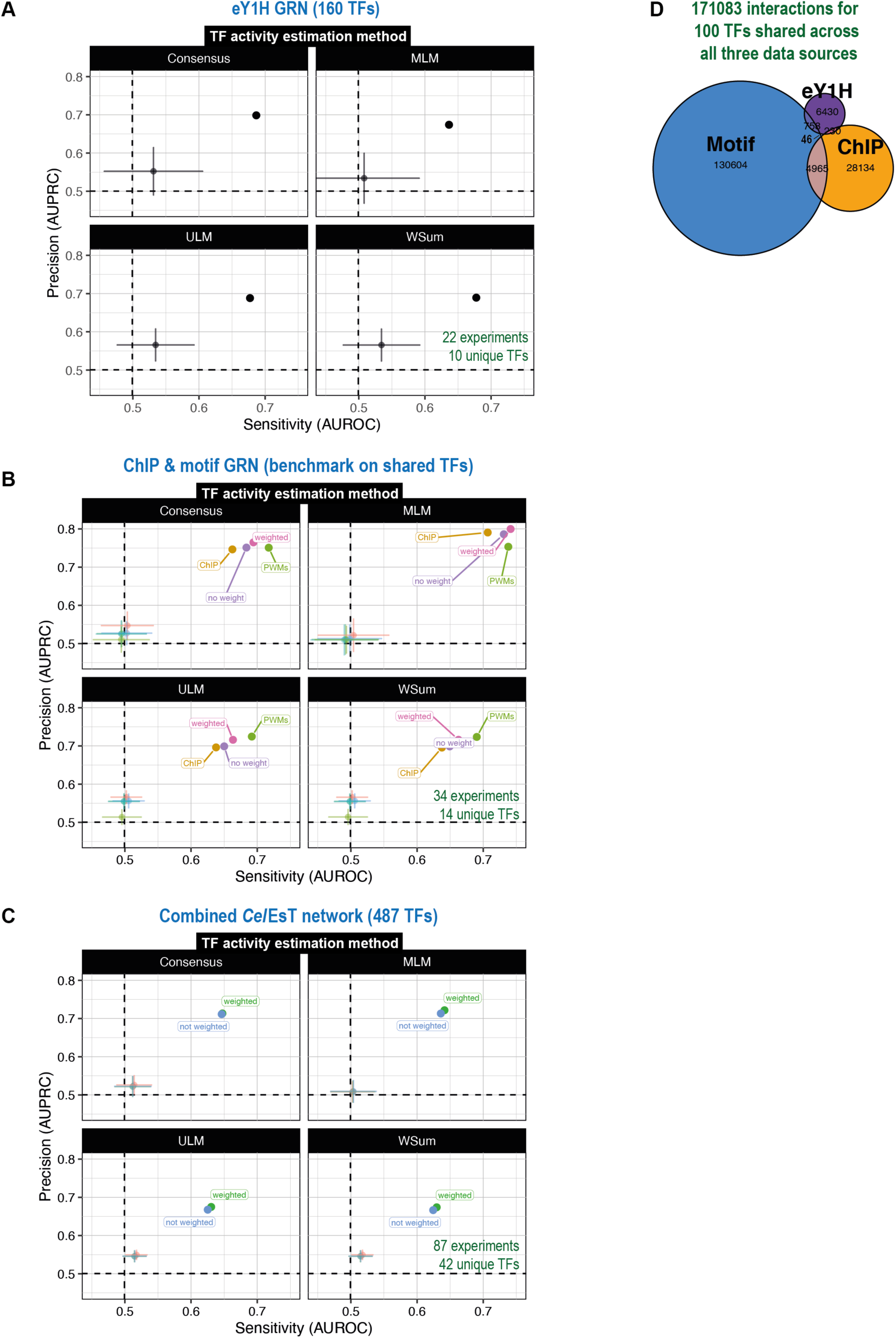
GRNs that combine targets from multiple data sources perform well despite few shared interactions (related to Fig 1) **A)** Benchmarking performance with different TF activity estimation methods (subpanels) for GRNs derived from TF-promoter interactions found in a large-scale enhanced yeast one-hybrid (eY1H) screen. Fig 1D shows benchmarking performance for multivariate linear model (‘mlm’) only. **B)** Benchmarking performance with different TF activity estimation methods (subpanels) for GRNs derived from ChIP-seq, DNA binding motif or a combination of the two with or without additional weight for shared targets. These GRNs are benchmarked on a common benchmarking set consisting of only TFs present in all networks to allow direct comparison. **C)** Benchmarking performance with different TF activity estimation methods (subpanels) for the *Cel*EsT network, which covers 487 TFs by combining data from multiple experimental sources. Fig 1E shows benchmarking performance for mlm only. **D)** Euler diagram showing contribution of TF-target interactions to the *Cel*EsT network from distinct experimental datasets only for 100 TFs common to all three. Total TF-target interactions shown above.

**Fig S4.**
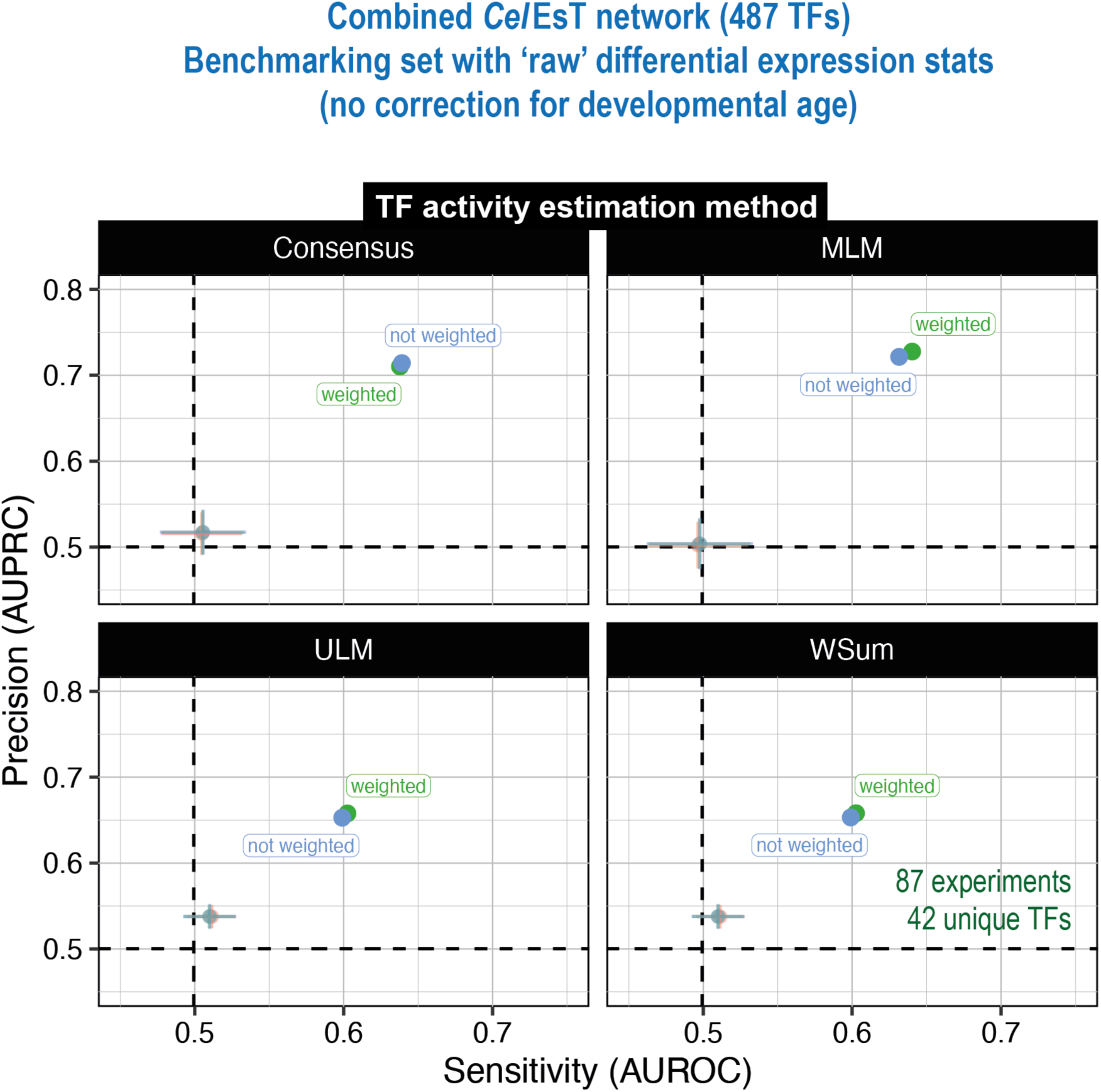
The combined *Cel*EsT combined network performs similarly well on a benchmarking dataset without developmental age correction. Benchmarking performance with different TF activity methods (subpanels) for the combined *Cel*EsT network (with or without additional weight for interactions shared across datasets) with a benchmarking set derived from differential expression analysis comparing treatment and control groups directly without accounting for any potential difference in developmental age.

**Fig S5.**
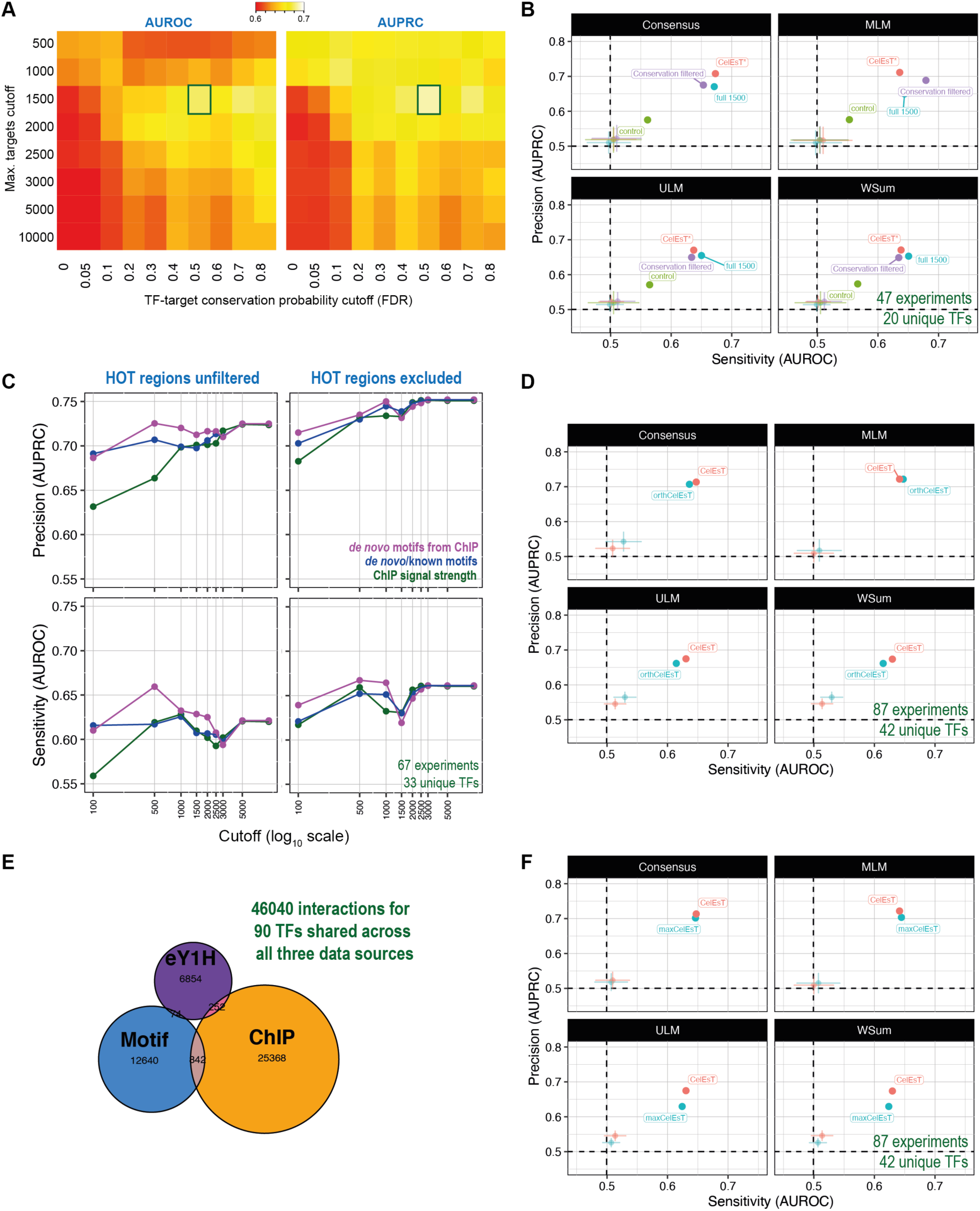
Motif conservation-based filtering of target genes to create GRNs with equal performance with fewer TF-target interactions (related to Fig 2) **A)** Heatmap shows benchmarking pipeline performance (AUROC left, AUPRC right) for motif-based GRNs with different initial elegans cut-offs and different cross-species conservation probability FDR cut-offs. The box shows the chosen network (top 1500 *elegans* targets filtered by FDR of 0.5) with the best performance for both AUROC and AUPRC. **B)** Benchmarking performance with different TF activity estimation methods (subpanels) for motif-based GRNs after conservation-based filtering (purple) versus the best-performing *elegans*-only motif-based GRN (‘full 1500’, blue), a control network with the same number of targets per TF selected from the best *elegans* motif matches (‘control’, green) and the *Cel*EsT network subsetted to the same TFs (‘CelEsT*’, red). Axes show performance (AUROC/AUPRC) in benchmarking pipeline. Transparent points show mean AUROC/AUPRC of randomly shuffled networks; error bars show standard deviation. The number of experiments and unique TFs in the benchmarking set overlapping with TFs in the GRN is noted on the panel in green text. Fig 2B shows benchmarking performance for ‘mlm’ only. **C)** Line shows evolution of AUPRC (above) and AUROC (below) with increasing cut-off for maximum number of targets per TF for GRNs derived from ChIP-seq data with (right) or without (left) exclusion of target genes within HOT regions. Colours indicate network with targets ordered by conservation probability from ChIP-derived *de novo* motifs (purple), known motifs from CisBP where applicable, else *de novo* motifs (blue) or ChIP peak signal strength (green). Note x-axis (cut-off) is on a log scale. Experiment/unique TF numbers in green text. Related to Fig 2D. **D)** Benchmarking performance with different TF activity estimation methods for the orth*Cel*EsT network. Details as for panel B. Fig 2D shows benchmarking performance for ‘mlm’ only. **E)** Euler plot shows TF-target interactions derived from each dataset in the orth*Cel*EsT network for those TFs shared between all three datasets. Related to Fig 2E. **F)** Benchmarking performance with different TF activity estimation methods for the max*Cel*EsT network. Details as for panel B.

**Fig S6.**
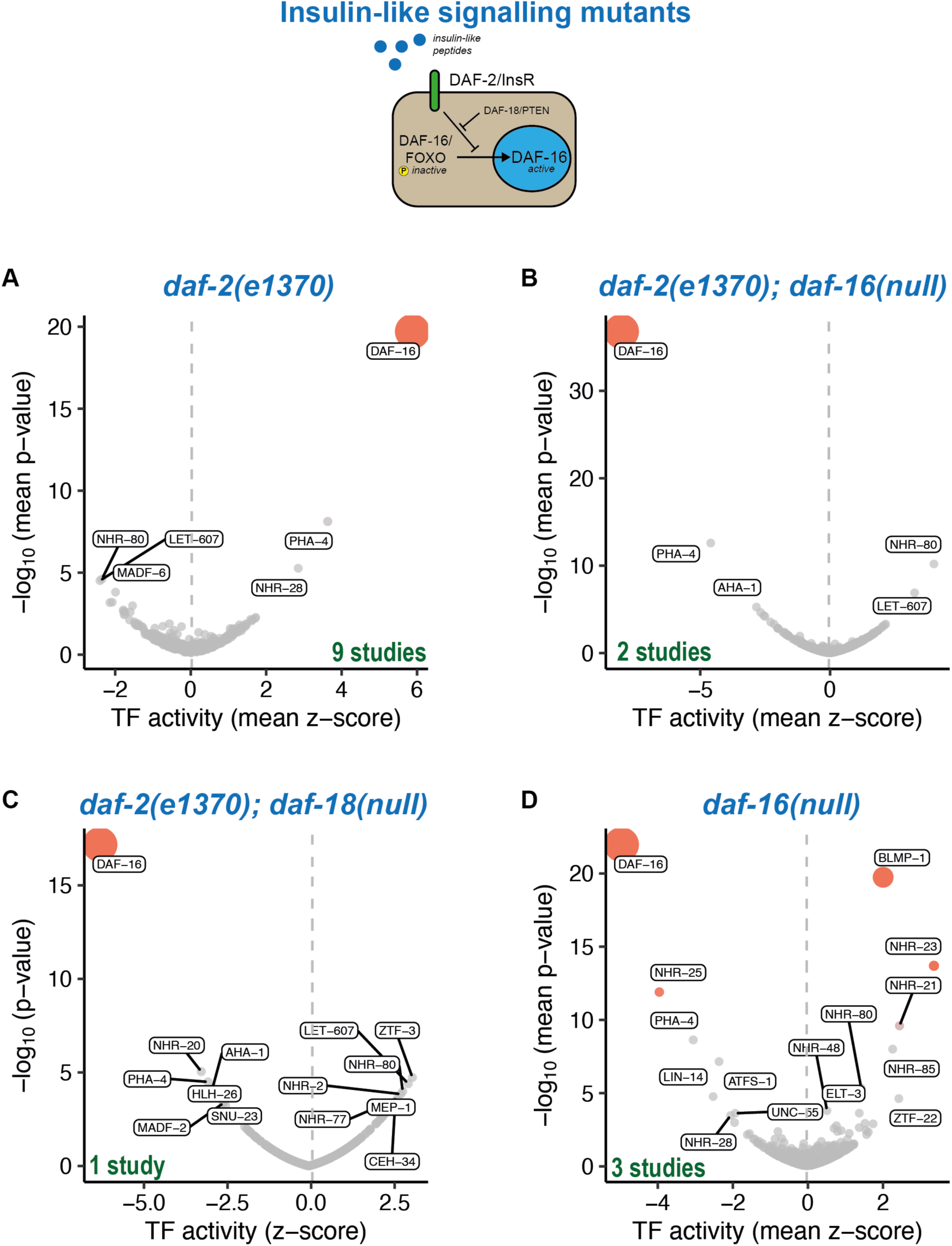
*Cel*EsT shows TF activity changes in mutants of multiple insulin/IGF-1-like signalling pathway components (related to Fig 3). Volcano plots shows mean TF activity z-score and geometric mean *p*-value. The bubble size is proportional to the *p*-value. **A)** 9 studies (see Fig S7A) comparing the severe *daf-2(e1370)* allele to wildtype controls. Note this panel is identical to that shown in Fig 3A and is provided here for ease of comparison. **B)** 2 studies (see Fig S7B) comparing *daf-2(e1370); daf-16(mu86)* to *daf-2(e1370)*. **C)** 1 study comparing *daf-2(e1370); daf-18(ok480)* to *daf-2(e1370)*. **D)** 3 studies (see Fig S7C) comparing *daf-16* null alleles to wildtype controls.

**Fig S7.**
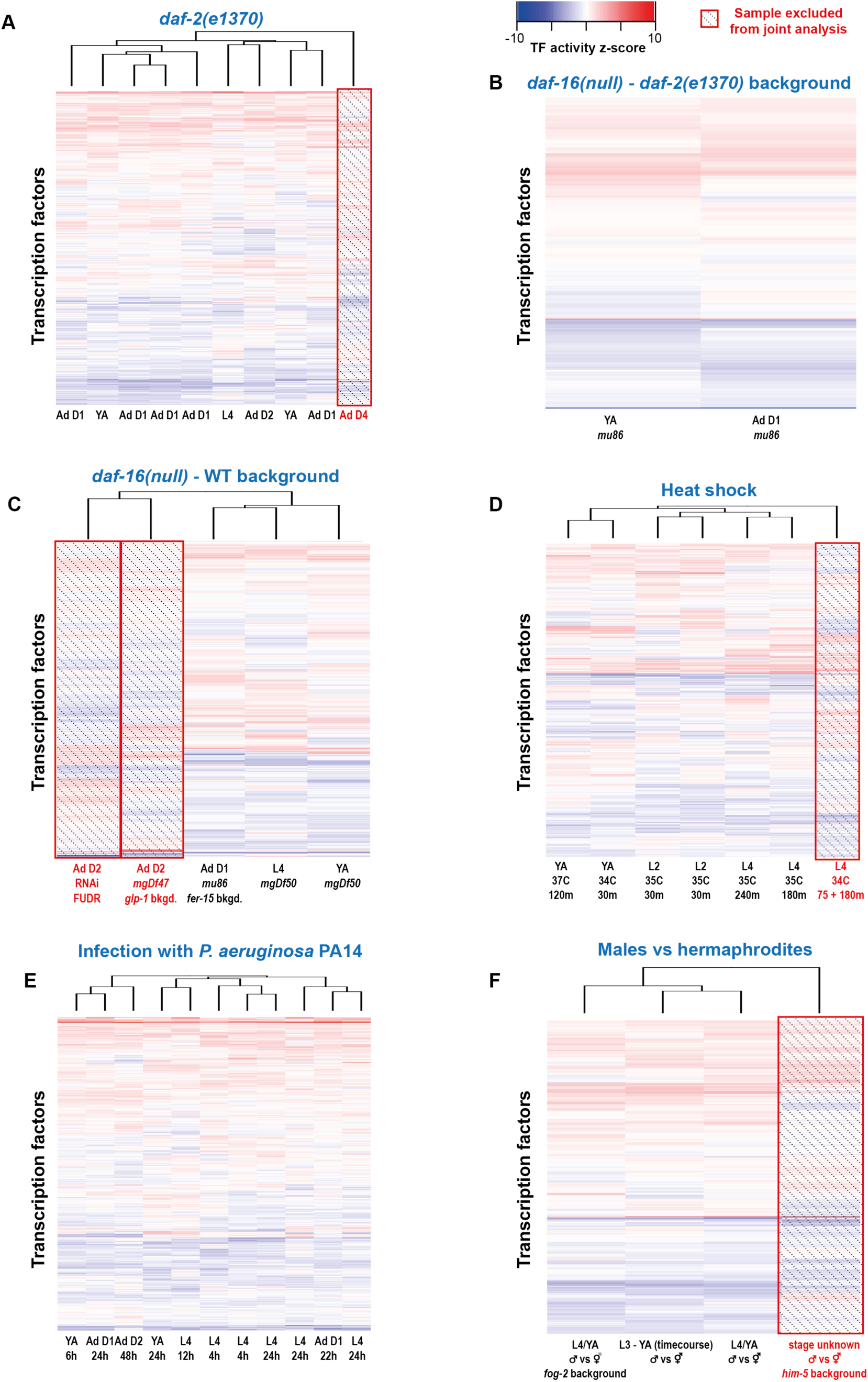
TF activity heatmaps for analyses combining results from multiple studies (related to Fig 3). Heatmaps show TF activity z-scores for each study considered for analysis for each condition. Study characteristics are annotated below; dendrograms above show study hierarchical clustering. Studies which were excluded as outliers and not included in the analyses shown in Fig 3 or Fig S6 are shaded out and marked with red text. **A)** 10 studies (1 excluded) comparing a severe mutant allele of the insulin receptor orthologue *daf-2* to wildtype controls. Related to Fig 3A, Fig S6A and Fig S8A. **B)** 2 studies comparing *daf-2(e1370); daf-16(mu86)* to *daf-2(e1370)*. Related to Fig S6B. **C)** 5 studies (2 excluded) comparing *daf-16(mu86)* animals to wildtype controls. Related to Fig S6D. **D)** 7 studies (1 excluded) comparing heat-shock treated animals to untreated controls. Related to Fig 3B and Fig S8B. **E)** 11 studies comparing animals infected with the pathogenic bacterium *P. aeruginosa* strain PA14 to unexposed controls. Related to Fig 3C and Fig S8C. **J)** 4 studies (1 excluded) comparing transcriptomes of male animals to hermaphrodites. Related to Fig 3D and Fig S8D.

**Fig S8.**
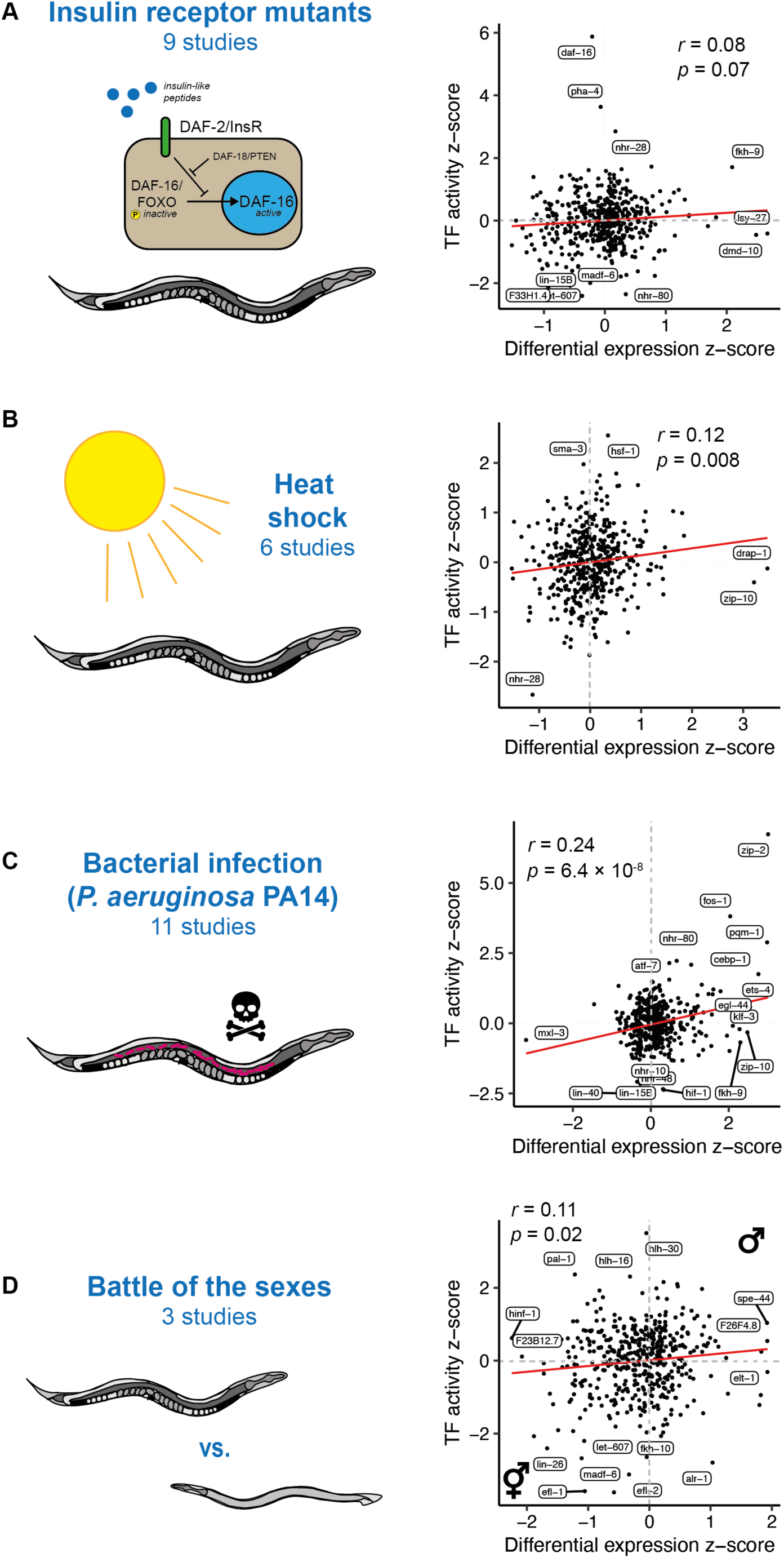
TF activity is independent of differential expression in many conditions but strongly related in bacterial infection (related to Fig 3). Scatterplots show mean differential expression z-scores for TF-encoding genes on the x-axis and *Cel*EsT-inferred activity for those TFs. Each plot is annotated with Pearson’s correlation coefficient (*r*) and *p*-value. All analyses controlled for developmental age during differential expression analysis, with DE stats then analysed using the multivariate linear model TF activity estimation method and the *Cel*EsT network. **A)** Analysis of 9 studies comparing a severe mutant allele of the insulin receptor orthologue *daf-2* to wildtype controls. See also Fig 3A and Fig S7A. **B)** Analysis of 6 studies comparing heat-shocked animals to untreated control animals. See also Fig 3B and S7D. **C)** Analysis of 11 studies comparing animals exposed to the pathogenic bacterium *Pseudomonas aeruginosa* PA14 to untreated controls. See also Fig 3C and S7E. **D)** Analysis of 3 studies comparing the transcriptome of male animals to that of hermaphrodites. See also Fig 3D and S7F.

**Fig S9.**
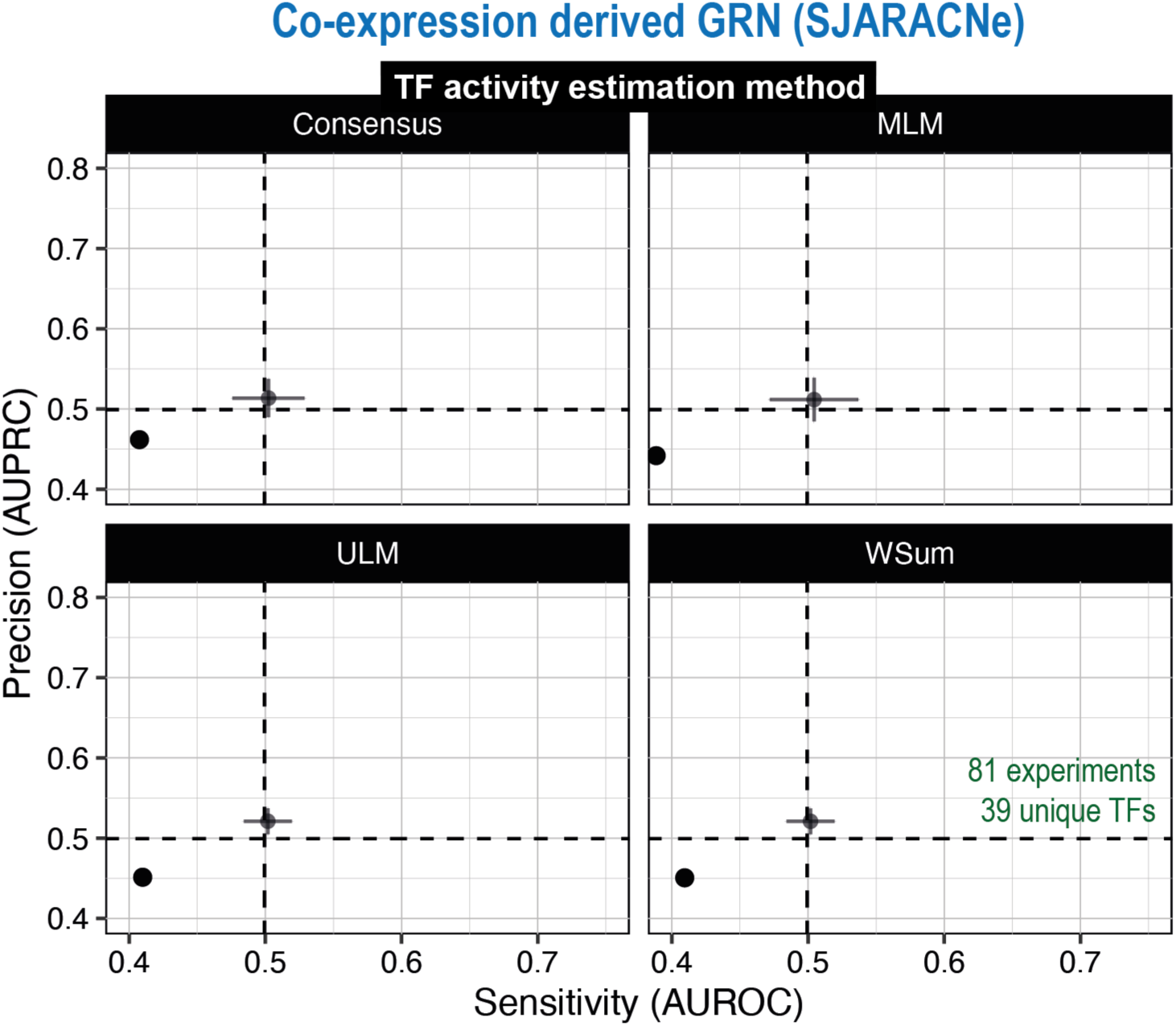
Co-expression derived gene regulatory network inferred by SJARACNe from a large RNA-seq dataset has no predictive capacity in benchmarking pipeline. TF activity estimation benchmarking performance for various methods for a GRN based on co-expression using edges computed by the SJARACNe algorithm using a large RNA-seq dataset from the CeNDR resource.

**Table S1.** TF annotation with modes of regulation

**Table S2.** Data sources for benchmarking dataset

**Table S3.** *Cel*EsT network and annotation

**Table S4.** orth*Cel*EsT network and annotation

**Table S5.** max*Cel*EsT network and annotation

**Table S6.** Datasets for *Cel*EsT validation

**Table S7.** Genome annotations for *Caenorhabditis* species

**File S1.** Customisable *R* script for performing TF activity estimation with *Cel*EsT

***Cel*EsT *R Shiny* app:** github.com/IBMB-MFP/CelEsT-app

