## Supplementary material for "*Cel*EsT: a unified gene regulatory network for estimating transcription factor activities in *C. elegans*": File S1

#start with empty workspace

rm(list = ls(all = TRUE))                                               

# turn off scientific notation for plots

options(scipen = 10000)

#### set working directory ####

# here create new folder and set working directory within it

dir.create("~/Desktop/CelEsT")
setwd("~/Desktop/CelEsT/")

dir.create("output")

#### DEFINE FUNCTIONS ####

# KUDOS to Kamil from biostars [https://www.biostars.org/p/171766/]
counts_to_tpm <- function(counts, featureLength, meanFragmentLength) {
  
  # Ensure valid arguments.
  stopifnot(length(featureLength) == nrow(counts))
  stopifnot(length(meanFragmentLength) == ncol(counts))
  
  # Compute effective lengths of features in each library.
  effLen <- do.call(cbind, lapply(1:ncol(counts), function(i) {
    featureLength - meanFragmentLength[i] + 1
  }))
  
  # Exclude genes with length less than the mean fragment length.
  idx <- apply(effLen, 1, function(x) min(x) > 1)
  temp_counts <- counts[idx,]
  temp_effLen <- effLen[idx,]
  temp_featureLength <- featureLength[idx]
  
  # Process one column at a time.
  tpm <- do.call(cbind, lapply(1:ncol(temp_counts), function(i) {
    rate = log(temp_counts[,i]) - log(temp_effLen[,i])
    denom = log(sum(exp(rate)))
    exp(rate - denom + log(1e6))
  }))
  
  # Copy the row and column names from the original matrix.
  colnames(tpm) <- colnames(temp_counts)
  rownames(tpm) <- rownames(temp_counts)
  
  return(tpm)
  
}

remove.duplicated.genes <- function(geneIDs){
  
  # here ensure no duplicated gene IDs in your data
  if(any(duplicated(geneIDs))){
    
    nmbr_dupl <- sum((duplicated(geneIDs)|duplicated(geneIDs, fromLast = TRUE)))
    
    uniquegeneids_fromDE_stats <- geneIDs[!(duplicated(geneIDs)|duplicated(geneIDs, fromLast = TRUE))]

    return(uniquegeneids_fromDE_stats)
    
    message(paste0(nmbr_dupl, " genes have been removed from analysis due to duplicated gene IDs"))
    
  } else {
    
    return(geneIDs)
    
  }
  
}

#### LOAD PACKAGES & FUNCTIONS ####

## First specify the packages of interest

packages <- c("stringr",
              "biomaRt",
              "openxlsx",
              "edgeR",
              "splines")

## Now load or install & load all packages
package.check <- lapply(packages, function(y){
  
  if (!require(y, character.only = TRUE)) {
    
    install.packages(y, dependencies = TRUE)
    
    library(y, character.only = TRUE)
    
  }
  
})

# Here define packages which need to be loaded through biocmanager

biocmanager_packages <- c("DESeq2",
                          "decoupleR") 

bioc_package.check <- lapply(
  
  biocmanager_packages,
  
  FUN = function(x) {
    
    if (!require(x, character.only = TRUE)) {
      
      if (!requireNamespace("BiocManager", quietly = TRUE)){
        
        install.packages("BiocManager")
        
      }
      
      BiocManager::install(x, dependencies = TRUE)
      
      library(x, character.only = TRUE)
      
    }
  }
)

# Here define packages to be loaded through Github

github_packages <- c("r-lib/conflicted",
                     "LBMC/RAPToR",
                     "LBMC/wormRef")

if (!requireNamespace("devtools", quietly = TRUE)){
  install.packages("devtools")}

gh_package.check <- lapply(
  github_packages,
  FUN = function(x) {
    if (!require(str_remove(x, ".*\\/"), character.only = TRUE)) {
      
      if (!requireNamespace("devtools", quietly = TRUE)){
        install.packages("devtools")}
      
      devtools::install_github(x, build_vignettes = TRUE)
      
      library(str_remove(x, ".*\\/"), character.only = TRUE)
      
    }
  }
)

#### INPUT DATA ####

parasite_mart <- useMart("parasite_mart", dataset = "wbps_gene", host = "https://parasite.wormbase.org", port = 443)

# set path to Table S1 (TF annotations) from manuscript supplementary files
TF_annotations <- read.xlsx("~/Downloads/Table_S1.txt")

# set path to Table S3 (annotated CelEsT network) from manuscript supplementary files
CelEsT_GRN <- read.table("~/Downloads/Table_S3.txt", 
                         sep = "\t",
                         header = TRUE)

## Here uncomment this code (RStudio shortcut Ctrl/Cmnd + Shift + C) and use it to upload your raw counts if needed - put in the address to your file
# 
# counts <- read.table(file = "~/path/to/file.txt", header = TRUE)

## Here uncomment this code (RStudio shortcut Ctrl/Cmnd + Shift + C) and use it to upload your existing DE stats if applicable
## If so, skip the next two sections of code ('DE no age correction' & 'DE with age correction')
# 
# DE_stats <- read.table(file = "~/path/to/file.txt",
#                        header = TRUE
#                        )

## You need to subset the DE_stats file to include only gene ID in first column and DE stats in second

# Choose your gene ID format to match your data

# see list of ID formats
print(head(wormRef::Cel_genes))

# set my ID format
myGeneIDformat <- "sequence_name"

# Below are two sections for performing DE analysis from raw counts. You can uncomment the code (shortcut Ctrl/Cmnd + Shift + C) that you need
# Both use DESeq2; if you use them please cite Love et al. 2014 doi.org/10.1186/s13059-014-0550-8
# If you use the analysis correcting for developmental age using RAPToR, please cite Bulteau et al. 2022 doi.org/10.1038/s41592-022-01450-0

# #### PERFORM DE ANALYSIS FROM RAW COUNTS ####
# 
# # If you wish to proceed from raw counts and not account for developmental age differences in your analysis,
# # use this section to derive your DE stats using DEseq2 as input to decoupler
# 
# # Enter your control and treatment sample IDs here. They should match the column names of your counts file
# 
# control_samples <- c("", "")
# treatment_samples <- c("", "")
#     
# # Here we ensure no duplicated gene IDs in your data
# gene_names_no_dups <- remove.duplicated.genes(counts$geneid)
# 
# counts_for_DE <- counts[counts[, 1] %in% gene_names_no_dups, ]
# row.names(counts_for_DE) <- gene_names_no_dups
#     
# counts_for_DE <- counts_for_DE[, c(control_samples, treatment_samples)]
# 
# tempcoldata <- data.frame(group = c(rep("control", times = length(control_samples)), rep("treatment", times = length(treatment_samples))))
# row.names(tempcoldata) <- c(control_samples, treatment_samples)
#     
# DErawdds <-  DESeqDataSetFromMatrix(countData = counts_for_DE,
#                                    colData = tempcoldata,
#                                    design = ~ group)
#     
# DErawdds <- DESeq(DErawdds)
#     
# DE_res <- as.data.frame(results(DErawdds))
# 
# DE_stats <- cbind(row.names(DE_res), DE_res[, "stat", drop = FALSE])

# #### PERFORM DE ANALYSIS WITH AGE CORRECTION WITH RAPToR ####
# 
# # Visualise age spans for developmental time course references to project samples onto
# plot_refs("wormRef")
# 
# # choose the appropriate reference below
# myRef <- "Cel_larv_YA"
# 
# # prepare reference 
# refdata <- prepare_refdata(myRef, 'wormRef', 5000)
# 
# control_samples <- c("", "")
# treatment_samples <- c("", "")
# 
# # here ensure no duplicated gene IDs in your data
# gene_names_no_dups <- remove.duplicated.genes(counts$geneid)
# 
# counts_for_DE <- counts[counts[, 1] %in% gene_names_no_dups, ]
# row.names(counts_for_DE) <- gene_names_no_dups
# 
# counts_for_DE <- counts_for_DE[, c(control_samples, treatment_samples)]
# 
# tempcoldata <- data.frame(group = c(rep("control", times = length(control_samples)), rep("treatment", times = length(treatment_samples))))
# row.names(tempcoldata) <- c(control_samples, treatment_samples)
#     
# # To estimate ages with RAPToR we need to convert the counts to TPM
# # we obtain the gene lengths from the RAPToR package and apply the counts_to_tpm function
# 
# # first we need to convert to WormBase Gene IDs
# 
# row.names(counts_for_DE) <- Cel_genes[match(row.names(counts_for_DE), Cel_genes[, myGeneIDformat]), "wb_id"]
# 
# my_temp_lengths <- sapply(row.names(counts_for_DE), function(x){
#   
#   mean(wormRef::Cel_genes[wormRef::Cel_genes$wb_id == x, "transcript_length"])
#   
#   })
#     
# my_temp_lengths <- my_temp_lengths[!is.na(my_temp_lengths)]
# 
# gene_there <- base::intersect(names(my_temp_lengths), row.names(counts_for_DE))
# 
# counts_for_DE_TPM <- counts_to_tpm(counts = counts_for_DE[gene_there, ],
#                                   featureLength = my_temp_lengths[gene_there],
#                                   meanFragmentLength = rep(100, times = ncol(counts_for_DE)))
#     
# # remove genes with zero expression across all samples
# counts_for_DE_TPM_expressed <- counts_for_DE_TPM[apply(counts_for_DE_TPM, 1, function(x){any(x != 0)}), ]
#     
# # transform for later use with RAPToR as per RAPToR vignette
# counts_for_DE_TPM_expressed_norm <- limma::normalizeBetweenArrays(counts_for_DE_TPM_expressed, method = "quantile")
# counts_for_DE_TPM_expressed_norm_log <- log1p(counts_for_DE_TPM_expressed_norm) # log1p(x) = log(x + 1)
# 
# # estimate sample ages    
# temp_sample_ae <- RAPToR::ae(samp = counts_for_DE_TPM_expressed_norm_log,                         # input gene expression matrix
#                                  refdata = refdata)
#     
# # Here we set the time buffer for the interpolated references on either side of the max/min sample ages
# # We set a buffer of one hour on each side
# # as reference times for embryo are in minutes, the offset will be 60 mins. Otherwise, times are in hours and the offset will be 1 hr
# 
# if(myRef == "embryo"){
#   
#   interpolation_offset <- 60
#   
# } else {
#   
#   interpolation_offset <- 1
#   
# }
# 
# # Here we set the age ranges for interpolated references
# interpolation_range <- range(temp_sample_ae$age.estimates[, 1]) + c(-interpolation_offset, interpolation_offset)
# 
# interpolation_index <- get_refTP(refdata, ae_values = interpolation_range)
# interpolation_index <- interpolation_index[1]:interpolation_index[2]
# 
# # Here we retreive interpolated counts from the reference dataset
# interpolated_ref_expression <- get_refTP(refdata,
#                                          ae_values = interpolation_range,
#                                          return.idx = FALSE)
# 
# interp_time <- refdata$time[interpolation_index]
# interp_tpm <- refdata$interpGE[, interpolation_index]
# 
# wormRef_gene_lengths <- sapply(row.names(interp_tpm), function(x){
#   
#   mean(wormRef::Cel_genes[wormRef::Cel_genes$wb_id == x, "transcript_length"])
#   
# })
# 
# # for use with DESeq, we need to convert the interpolated TPM values back to counts
# # we arbitrarily set a library size of 25 million reads for converting the reference samples into hypothetical counts
# libsize <- 25e6
# 
# interp_count <- t((t (exp(interp_tpm) - 1)/ 1e6) * (libsize / median(wormRef_gene_lengths))) * wormRef_gene_lengths
# interp_count[interp_count < 0] <- 0
# interp_count <- round(interp_count)
# 
# # now we have the age estimates. We do GLM with edgeR with glmFit. 
# # We include batch (sample vs reference data), variable of interest (mutation, group reference with control) and developmental time modelled with splines ( )
# # First we need to determine optimal number of spline df by fitting different models
# 
# # We apply models with various degrees of freedom to find a good number for the modelling that maximises R-sq with a low df
# 
# df_SSQ <- sapply(1:8, function(df_param){
#   
#   sum(residuals(lm(t(interp_tpm) ~ splines::ns(interp_time, df = df_param))) ^2)
#   
# })
# 
# # here find the point at which the curve plateaus appropriately
# # The heuristic threshold is arbitrarily set at 0.01
# threshold <- 0.01
# diff_1 <- diff(df_SSQ)
# 
# temp_plateau <- which.max((diff_1 / diff_1[1]) < threshold) 
# 
# # Having found an appropriate plateau, proceed to fit model with appropriate spline df 
# 
# # restrict to genes with at least 5 counts in 1 sample
# counts_forDE <- counts_for_DE[apply(counts_for_DE, 1, max) > 5, ]
# 
# # combine counts for samples and interpolated references
# combine_count <- do.call(cbind, format_to_ref(counts_for_DE, interp_count)[1:2])
# 
# combine_coldata <- data.frame(time = c(temp_sample_ae$age.estimates[, 1], interp_time),
#                               strain = c(tempcoldata[row.names(temp_sample_ae$age.estimates), 1], rep("control", ncol(interp_count))),
#                               batch = rep(c("sample", "reference"), c(ncol(counts_forDE), ncol(interp_count))))
# 
# combine_coldata$strain <- factor(combine_coldata$strain, levels = c("control", "treatment"))
# combine_coldata$batch <- factor(combine_coldata$batch, levels = c("sample", "reference"))
# 
# row.names(combine_coldata) <- c(row.names(temp_sample_ae$age.estimates), colnames(interp_count))
# 
# # We estimate gene dispersions on sample data alone
# 
# dd0 <- DESeqDataSetFromMatrix(countData = combine_count[, 1:length(c(control_samples, treatment_samples))],
#                               colData = combine_coldata[1:length(c(control_samples, treatment_samples)), ],
#                               design = ~ strain)
# 
# dd0 <- estimateSizeFactors(dd0)
# dd0 <- estimateDispersions(dd0, fitType = "local") 
# d0 <- dispersions(dd0) # store dispersions
# d0[is.na(d0)] <- 0 # remove NAs
# 
# # We set the formula for use in the DE analysis
# formula <- paste0("~ splines::ns(time, df = ", temp_plateau, ") + batch + strain")
# 
# # combine data for final DE analysis
# dd1 <-  DESeqDataSetFromMatrix(
#   countData = combine_count,
#   colData = combine_coldata,
#   design = as.formula(formula) # use df found above
# )
# 
# dd1 <- estimateSizeFactors(dd1)
# 
# # inject dispersions from sample-only model
# dispersions(dd1) <- d0
# 
# dd1 <- nbinomWaldTest(dd1)
# 
# DE_res <- as.data.frame(results(dd1, contrast = c("strain", "treatment", "control")))
# 
# # change ID format back to original format
# row.names(DE_res) <- Cel_genes[match(row.names(DE_res), Cel_genes$wb_id), myGeneIDformat]
# 
# DE_stats <- cbind(row.names(DE_res), DE_res[, "stat", drop = FALSE])

#### FORMAT DIFFERENTIAL EXPRESSION STATS FOR DECOUPLER ####

# Check gene IDs are ok - we want sequence names to match CelEsT

# the str_remove() expression removes the letter corresponding to transcript isoform to convert transcript IDs into gseq names
DE_stats <- DE_stats[str_remove(DE_stats[, 1], "\\.[a-z]{1,2}") %in% unlist(Cel_genes), ]

DE_stats_genes <- remove.duplicated.genes(DE_stats[, 1])
DE_stats <- DE_stats[match(DE_stats_genes, DE_stats[, 1]), ]

DE_stats <- DE_stats[, 2, drop = FALSE]

row.names(DE_stats) <- DE_stats_genes

# replace NAs with 0s
DE_stats[is.na(DE_stats)] <- 0

# convert to sequence names if applicable
row.names(DE_stats) <- Cel_genes[match(row.names(DE_stats), Cel_genes[, myGeneIDformat]), "sequence_name"]

#### ESTIMATE TF ACTIVITY WITH DECOUPLER ####

# Please cite the CelEsT paper (Perez 2024) and the decoupler paper (Badia-i-Mompel et al. 2022 doi.org/10.1093/bioadv/vbac016)

# Here we perform the TF activity estimation with the multivariate linear model method (recommended)
# To use other methods, change the argument to statistics or supply multiple methods as a vector (e.g. statistics = c("mlm", "ulm"))
# Estimation can take a couple of minutes. mlm is the fastest method, other methods are considerably slower
DEdata_decouple <- decoupleR::decouple(
  mat = DE_stats[, 1, drop = FALSE], 
  network = CelEsT_GRN,
  .source = "source",
  .target = "target",
  statistics = "mlm",
  args = list(mlm = list(.mor = "weight")),
  consensus_score = FALSE
)

# annotate output with additional TF identifiers

DEdata_decouple <- cbind(DEdata_decouple,
                         Cel_genes[match(DEdata_decouple$source, Cel_genes$sequence_name), c("wb_id", "public_name")])

# capitalise TF names
DEdata_decouple[, "public_name"] <- toupper(DEdata_decouple[, "public_name"])

# order by p-value
DEdata_decouple <- DEdata_decouple[order(DEdata_decouple$p_value), ]

# annotate according to MOR from UniProt/WormBase
DEdata_decouple <- cbind(DEdata_decouple, TF_annotations[match(DEdata_decouple$source, TF_annotations$wormbase_gseq), c("Wormbase_mode_of_regulation", "Uniprot_mode_of_regulation")])

DEdata_decouple[, c("Wormbase_mode_of_regulation", "Uniprot_mode_of_regulation")] <- str_replace_all(unlist(DEdata_decouple[, c("Wormbase_mode_of_regulation", "Uniprot_mode_of_regulation")]), "-1", "REPRESSOR")
DEdata_decouple[, c("Wormbase_mode_of_regulation", "Uniprot_mode_of_regulation")] <- str_replace_all(unlist(DEdata_decouple[, c("Wormbase_mode_of_regulation", "Uniprot_mode_of_regulation")]), "0", "BIFUNCTIONAL")
DEdata_decouple[, c("Wormbase_mode_of_regulation", "Uniprot_mode_of_regulation")] <- str_replace_all(unlist(DEdata_decouple[, c("Wormbase_mode_of_regulation", "Uniprot_mode_of_regulation")]), "1", "ACTIVATOR")

# output results to file
write.table(DEdata_decouple,
            "output/TF_activity_estimations.txt",
            col.names = TRUE,
            row.names = FALSE)
